# Early multi-omic signatures and machine learning models predict cardiomyocyte differentiation efficiency and enable robust hPSC differentiation to cardiomyocytes

**DOI:** 10.64898/2026.01.08.698025

**Authors:** Austin K. Feeney, Aaron D. Simmons, Elizabeth F. Bayne, Yanlong Zhu, Mason R. Pentes, Paulo F. Cobra, Jianhua Zhang, Timothy J. Kamp, Ying Ge, Sean P. Palecek

## Abstract

Protocols for generating cardiomyocytes (CMs) from human pluripotent stem cells (hPSCs) have existed for nearly two decades, yet manufacturing variability in terminal cell identity continues to limit clinical translation. To uncover the origin of fate divergence during hPSC-CM differentiation, we performed temporal transcriptomics, proteomics, and metabolomics of high and low efficiency differentiations. We identified significant early multi-omic divergence between differentiation batches and key pathways underlying fate divergence at critical differentiation stages included Wnt, MAPK, and glucose metabolism. Machine learning models trained on early candidate gene markers predicted hPSC-CM purity better than models using canonical cardiac development markers. Lastly, multi-omic insights informed perturbations, including Wnt and MAPK inhibition, which produced higher CM purities and yields. Our results showcase multi-omic analysis coupled with machine learning models as a powerful tool to identify cell fate determinants and enable robust manufacturing of complex cell products such as hPSC-derived cell therapies.

**Teaser:** Multi-omic analysis of hPSC-CM differentiation efficiency reveals early predictive features and enables robust differentiation.

## Introduction

The last decade has been marked by significant progress in using cell and gene therapies to treat human disease. In 2017, the FDA approved the first chimeric antigen receptor T cell (CAR T) therapy for acute lymphoblastic leukemia accelerating efforts to establish automated, scalable cell culture practices^1,2^. Human pluripotent stem cells (hPSCs) offer unmatched therapeutic potential due to their unique ability to generate any cell type in the body^3^. However, the multipotent nature of hPSCs presents a major challenge in cell manufacturing to reproducibly generate the desired cell type with minimal contamination by other undesired cell lineages. Thus, there is an urgent need to develop approaches to monitor differentiation progression and feedback-controlled closed-loop manufacturing processes to unlock the potential of hPSC-derived cell therapies.

The ability to consistently manufacture a safe and efficacious hPSC-cardiomyocyte (hPSC-CM) cell product is critical because ischemic heart disease remains the leading global cause of death^4^. hPSC-CMs have demonstrated immense potential in developmental biology research^5,6^, drug screening^7,8^, disease modeling^9,10^, and cell therapy^11,12^. However, broad application of hPSC-CMs remains limited by high variability across differentiation protocols, cell lines, and batches. The well-established small molecule-based GiWi (GSK3β inhibitor/Wnt inhibitor) protocol modulates Wnt signaling through sequential addition of CHIR99021 (CHIR) for mesoderm induction and IWP2 for cardiac specification^13,14^, whereas growth factor-based protocols typically employ a combination of BMP4, FGF2, and Activin A^15,16^. Both hPSC-CM differentiation paradigms are highly sensitive to cell density, signaling molecule concentrations, timing, and other factors, leading to terminal CM purities ranging from complete failure to >95%^17,18^. Moreover, the onset of lineage divergence can vary widely, resulting in a variety of off-target cell types during and at the end of differentiation^19–21^. However, early predictive markers capable of forecasting hPSC-CM differentiation outcome remain limited, preventing feedback-controlled interventions to rescue CM fate.

Although prior studies have advanced understanding of stage-specific signaling and marker acquisition during hPSC-CM differentiation, most have relied on limited marker analyses, limited sampling stages, and high or unreported differentiation purities. These experimental designs have hindered the ability to understand differences between failed and successful differentiations^22–24^. While transcriptomic^25^, proteomic^26^, metabolomic^27–29^, and dual transcriptomic and proteomic^30,31^ approaches have been utilized, metabolomic studies remain scarce despite the central role of metabolic transitions during *in vivo* cardiomyocyte development^32–34^.

To address these gaps, we employed a unique multi-omic discovery approach integrating transcriptomics, proteomics, intracellular metabolomics, and extracellular metabolomics across the differentiation time course of high and low efficiency differentiations. Our goals were to 1) define molecular and functional mechanisms of hPSC-CM fate commitment, 2) identify early predictive markers of terminal hPSC-CM purity, and 3) inform strategies to rescue low purity hPSC-CM differentiation. Using a two-factor experimental design with time-resolved multi-omic sampling of high and low efficiency purity differentiations, we decoupled temporal signaling dynamics common to generic differentiation of hPSCs in response to cardiogenic signaling cues from the developmental processes unique to high purity CM fate commitment *in vitro*. Notably, we discovered that early multi-omic divergence predicts hPSC-CM differentiation outcome. Moreover, early differences in gene expression between high and low efficiency hPSC-CM differentiations informed machine learning models that outperformed canonical cardiac differentiation genes in predicting terminal CM purity. Furthermore, multi-omic pathway findings enabled the identification of differential metabolic functional phenotypes between high and low efficiency hPSC-CM differentiations and informed a small molecule-based Wnt and MAPK inhibition approach to improve hPSC-CM differentiation purity.

## Results

### Early multi-omic divergence predicts hPSC-CM differentiation outcome

As a first step toward identifying multi-omic markers and mechanisms underlying divergent cell fate outcomes during hPSC-CM differentiation, we collected parallel samples throughout the differentiation of hPSCs to hPSC-CMs via the GiWi protocol for multiple batches that generated different CM purities. At each collection timepoint (Day 0, 2, 4, 6, 8, 10, 13, and 16; i.e. the days of media replacement), 3-4 wells were harvested for paired collections of media for extracellular metabolite analysis and cells for analysis of intracellular metabolites and proteins. An additional 3-4 wells were collected for RNA analysis (**Figure 1A**). At the end of the differentiation, parallel wells from each batch were assessed for CM purity via flow cytometry for cardiac troponin T (cTnT). These terminal CM percentages were used to classify batches as high or low efficiency CM differentiations (**Figure 1B, Figure S1A-B**). The selected batches for multi-omic analysis had significantly different terminal CM purities of approximately 25% (low efficiency, magenta) and 75% (high efficiency, turquoise). To validate the quality of the multi-omic data, we assessed the relative protein (**Figure 1C**) and transcript (**Figure 1D**) abundances of cTnT/*TNNT2* at the end of the differentiation and confirmed that protein and mRNA measurements both exhibited a significant enrichment of approximately three-fold in high versus low efficiency batches, in concordance with the flow cytometry and immunocytochemistry (ICC) of cTnT (**Figure 1E**).

**Figure 1.**
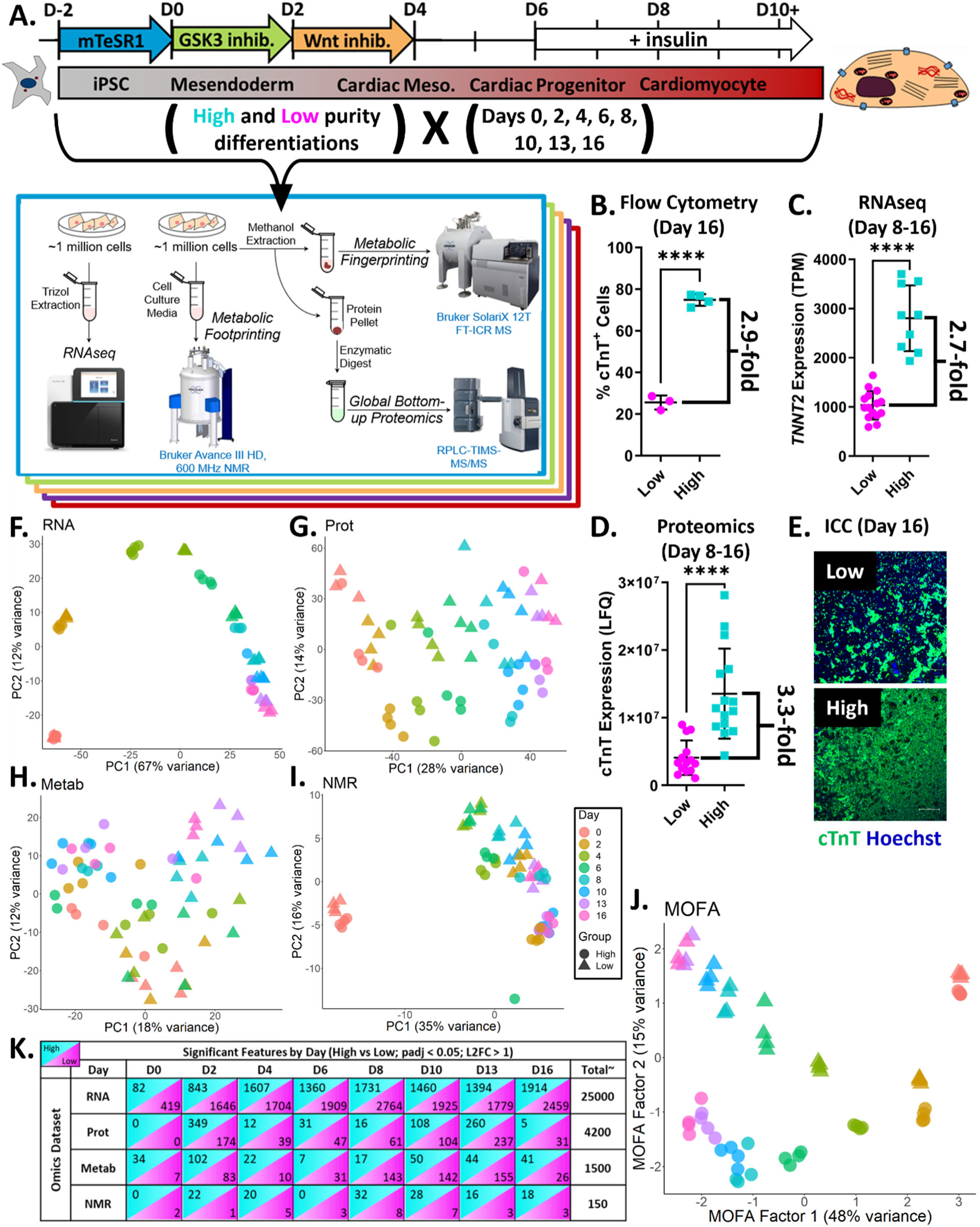
Early multi-omic divergence predicts hPSC-CM differentiation outcome. **A)** Schematic of the hPSC-CM differentiation protocol (WTC11 hiPSCs) and the two-factor sampling design (differentiation efficiency and timepoints). **B)** Terminal percentage of cTnT+ cells (% CMs) via flow cytometry used to assign high and low efficiency batches (n = 3-4 well replicates from one differentiation per group). **C)** RNA TPM of *TNNT2* for samples collected between Days 8-16 for high and low efficiency batches (n = 3-4 well replicates per group from one differentiation, except for Days 10 and 13 in high efficiency group where n = 1 and 2 respectively) **D)** Proteomic abundance (LFQ) of cTnT for samples collected between Days 8-16 for high and low efficiency batches (n = 3-4 well replicates per group from one differentiation). **E)** Representative immunocytochemistry (ICC) images of cTnT expression (Scale bars = 500µm). **F-I)** Principal component analyses (PCA) of the **F)** transcriptome (RNA), **G)** proteome (Prot), **H)** intracellular metabolome (Metab), and **I)** extracellular metabolome (NMR) (color encodes sample timepoint, shape encodes terminal purity). **J)** Multi-omic factor analysis (MOFA) plot of factors 1 and 2 incorporating all -omes. Multi-omic data were collected from WTC11 hiPSCs. **K)** Differential feature abundance at each timepoint for each -ome. P-values from unpaired t-tests. All data are represented as mean ± SD. TPM = transcripts per million. LFQ = label free quantitation.

To provide a high-level, unbiased view of the multi-omic data for high and low efficiency CM batches, we generated PCA plots of all features identified in each individual-ome for the entire dataset (**Figure 1F-I**). Interestingly, each -ome exhibited unique temporal trends for high- and low-efficiency batches and demonstrated the potential to distinguish between these divergent outcomes even at early timepoints. Next, we performed multi-omic factor analysis (MOFA) to identify multi-omic trajectories of each batch^35^. Through this analysis, we identified two multi-omic factors responsible for a majority of the variance among the data (**Figure 1J, Figure S1C-E**). Factor 1 correlated to the time course of the differentiation cascade, with higher positive values corresponding to early timepoints, and lower or negative values corresponding to late timepoints. Accordingly, Factor 1 values correlated with features enriched at later timepoints (myosins, troponins, extracellular matrix (ECM) proteins, and carnitines; **Figure S1F-I**). Interestingly, the low efficiency group exhibited more negative Factor 1 values than the high efficiency group at all timepoints except Day 0, suggesting it progressed through differentiation at a faster rate than the high efficiency group. Factor 2 encoded much of the separation between the high and low purity batches, with the low purity samples exhibiting higher or more positive factor values than the high purity samples at every timepoint. Further investigation of these factors identified that Factor 1 was driven predominantly by RNA features, with moderate contributions from protein features (**Figure S1E**). In contrast, Factor 2 was evenly weighted as its strongest positive drivers (i.e. those enriched in low efficiency batches) included ECM and ECM-related genes and proteins (e.g. COL5A1/2, COL1A1, DCN, LEPREL2, *ADAMTS5, LUM, POSTN*), lipid or lipid-associated features (e.g. ANXA1, C21H31N9O5, C22H35N3O5), and calcium-related features (e.g. *HPCAL4, SLC24A3, S100A10*) (**Figure S1E**, **J-M**). The top negative-weighted features (i.e. those enriched in high efficiency batches) included transcription factors/nuclear effectors (SALL4, MUSTN1*, TBX3, IRX3, HHEX)*, transmembrane receptors (*CXCR4, APLNR*), and bioactive metabolites (2-3,4dimethylphenyl-ethylbeta-D-glucopyranoside, 12:4_3 fatty acyl hexoside, tetradecanoyl-L-carnitine).

Lastly, we performed differential analysis between high and low efficiency batches for each -ome at each timepoint (**Figure 1K**). Notably, significant divergence between high and low efficiency batches was evident as early as Day 2, generally increased with time, and was apparent across all -omes. With these promising results identifying the potential to distinguish different fate outcomes very early in differentiation, we extended our analysis to include additional transcriptomic samples for additional cell lines and batches focused on assessing cell fate divergence at early timepoints, up to the CPC stage.

### Early divergence between high and low efficiency hPSC-CM differentiations is conserved across cell lines

To provide more robust insight into early cell fate divergence in hPSC-CM differentiation, we extended the transcriptomic analysis to include 6 additional batches from 3 cell lines (H9 hESCs, IMR90-4 hiPSCs, and WTC11 hiPSCs; one high and one low efficiency batch each). Due to timing differences in the optimal differentiation protocols among these cell lines, we refer to differentiation steps where key differentiation modulators are added or removed in each protocol as opposed to sample timepoints^13,36^. Using this terminology, the +CHIR differentiation step corresponds to Day 0 for both protocols, the +IWP2 differentiation step corresponds to Day 2 for the WTC11 and Day 3 for H9 and IMR90-4 (Day 2/3), the −IWP2 differentiation step corresponds to Day 4/5, and the −INS differentiation step corresponds to Day 6/7 (**Figure 2A**). We selected high and low efficiency hPSC-CM differentiation batches for each cell line based on terminal flow cytometry assessment for transcriptomic analysis (**Figure 2B**). For these early differentiation steps, the transcriptomic data was combined with the data from WTC11 hiPSC samples shown in **Figure 1** and batch-corrected prior to downstream analyses.

**Figure 2.**
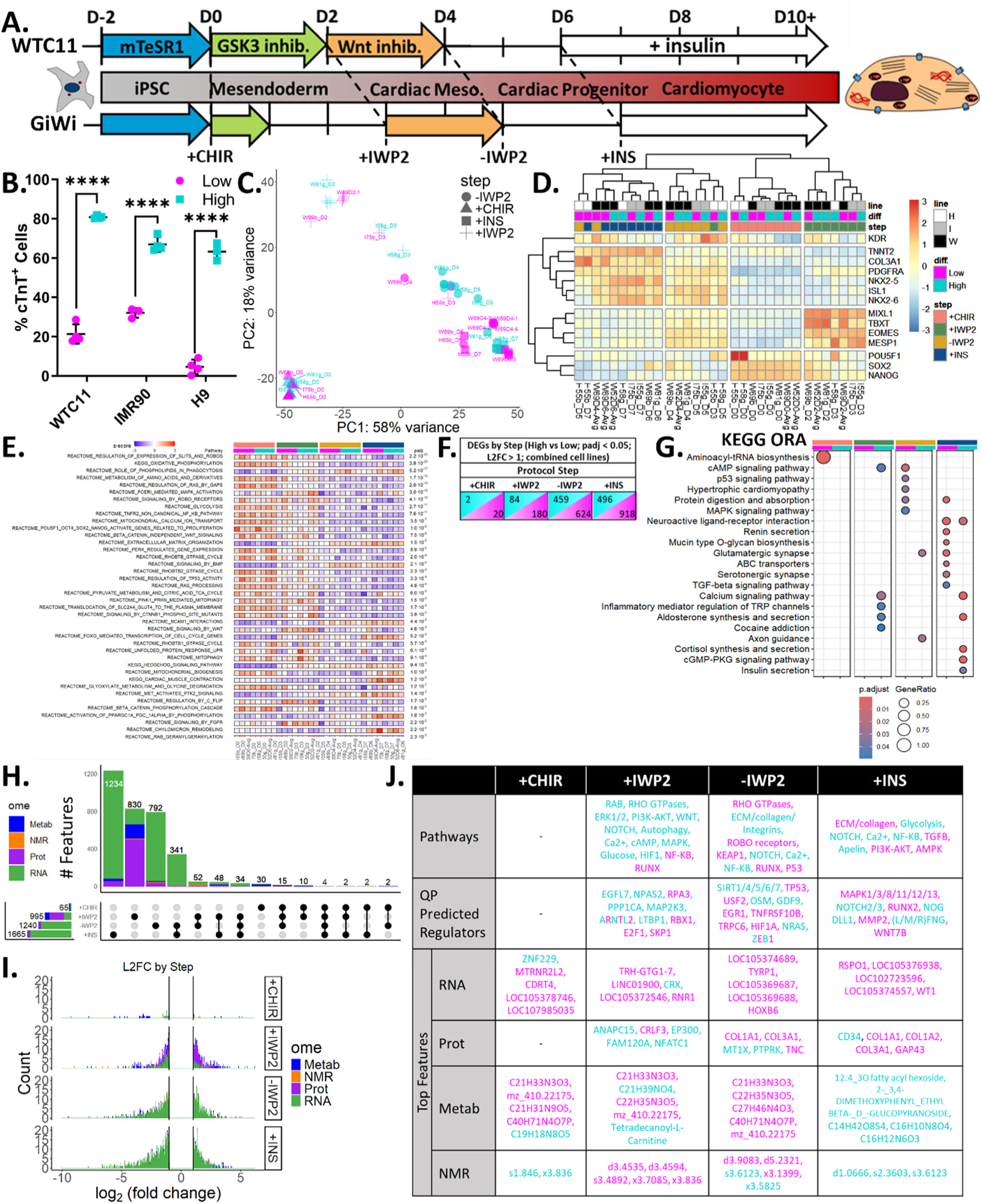
Early divergence between high and low efficiency hPSC-CM differentiations is conserved across cell lines. **A)** Schematic of the hPSC-CM differentiation protocol for WTC11 hiPSCs and the GiWi cell lines (IMR90-4 hiPSCs and H9 hESCs). The +IWP2, −IWP2, and +INS differentiation steps require differential timing per the optimized protocol across cell lines. **B)** Terminal percentage of cTnT+ cells (% CMs) via flow cytometry used to assign high and low efficiency batches (n = 3-4 well replicates for one differentiation per group). **C)** PCA of the batch-corrected transcriptomic data from these samples and Figure 1 samples. **D)** Transcriptomic heatmap (TPM) of canonical stage-specific gene markers throughout the hPSC-CM differentiation. **E)** GESECA heatmap of Reactome and KEGG terms enriched at each differentiation step. **F)** Table of differentially expressed genes at each differentiation step from the batch-corrected transcriptomic data. **G)** Overrepresentation analysis (ORA) of KEGG pathways enriched in the high or low efficiency groups at each differentiation step. **H)** UpSet plot of differential features at each differentiation step. **I)** Histogram for the number of differentially abundant -omic features with the corresponding log2 fold change between high and low efficiency differentiation groups at each differentiation step. **J)** Top differential pathways, predicted regulators, and features for each -ome at each differentiation step; color-coded by direction of change (turquoise = enriched in high efficiency, magenta = enriched in low efficiency). P-values from unpaired t-tests. All data are represented as mean ± SD. TPM = transcripts per million. Line = cell line (H = H9, I = IMR90-4, W = WTC11).

Similarly to the PCA and differential expression analysis in **Figure 1F,J**, we observed PC1 encoding a temporal axis, with significant divergence between high and low purity batches as early as the second step in the differentiation protocol (+IWP2, **Figure 2C**). Moreover, there was a high degree of sample clustering in the PCA between high and low purity batches at the +CHIR differentiation step, which demonstrated that the divergence between high and low efficiency groups at the +IWP2 differentiation step was not simply driven by poor quality hPSCs corresponding to Day 0. Notably, timepoint alignment by differentiation step (+CHIR, +IWP2, −IWP2, +INS) was confirmed by clear sample clustering according to differentiation stage across all cell lines and purities using canonical hPSC-CM trajectory marker genes (**Figure 2D**). This included high expression of undifferentiated state markers *POU5F1*, *SOX2*, and *NANOG* at the +CHIR differentiation step and high expression of mesoderm state markers *MIX1L*, *TBXT*, *EOMES*, and *MESP1* at the +IWP2 differentiation step. At the −IWP2 and +INS differentiation steps, CPC stage markers were highly expressed with increased expression of the cardiomyocyte marker, *TNNT2,* and the off-target cardiac fibroblast marker, *COL3A1,* at the +INS differentiation step. Interestingly, several low purity samples clustered with high purity samples at a later differentiation step (**Figure 2D**), again suggesting that low purity differentiation batches may progress through differentiation at a faster rate than the high purity batches.

To further investigate differences in differentiation rate and off-target fates, we utilized Platform-agnostic CellNet (PACNet) analysis to investigate acquisition of cell and tissue type-specific gene signatures^37^. Notably, low purity hPSC-CM progenitors displayed a significantly increased heart classification score in comparison to their high purity counterparts early in differentiation (-IWP2 step, D4). However, this relationship switched later in differentiation (D10-D15), with high purity hPSC-CM samples exhibiting significantly higher heart classification scores than low purity samples (**Figure S2A**). Moreover, low purity hPSC-CM progenitors lost their hPSC signature significantly faster than high purity hPSC-CM progenitors, with decreased hPSC classification scores early in differentiation (D4-D8) in comparison to their high efficiency counterparts (**Figure S2B**). Low purity hPSC-CM progenitors exhibited significantly increased off-target signatures, including fibroblast and endoderm (combination of lung, liver, and intestine classification scores) classification scores, both early (-IWP2, D4) and at the end of hPSC-CM differentiation (**Figure S2C-D**).

Across the combined, two-factor transcriptomic dataset, we performed a multi-conditional gene set enrichment analysis (GSEA) technique called gene set co-regulation analysis (GESECA) to identify gene sets with high internal gene correlation representing coordinated and significant changes in cellular functions across the dataset (**Figure 2E**)^38^. The top identified gene sets were dominated by temporal differences conserved between the high and low efficiency groups (i.e. enriched at the same differentiation step across both groups). This demonstrated that among the top gene sets, few exhibited potential to discriminate between high and low efficiency differentiation batches because the gene sets with the highest degree of variation were common to hPSC-CM differentiation stages regardless of differentiation purity. Across high and low efficiency differentiation groups, we found that the +CHIR step was enriched in numerous terms related to “proliferation”, “RHOBTB”, “glycolysis” and “central energy metabolism”, “mitochondrial biogenesis”, and “Wnt/beta-catenin regulation”. The +IWP2 stage was enriched for terms related to “MAPK”, “Wnt”, “c-FLIP”, and “FGFR”. The −IWP2 stage was enriched for terms relating to “phospholipids”, “ECM”, “BMP”, “NCAM1”, and “PTK2” and these continued through the +INS stage. Finally, several pathways enriched specifically at the +INS stage included “PGC1A”, “cardiac muscle contraction”, “chylomicron remodeling”, and “FOXO” in addition to the re-enrichment of “mitochondrial biogenesis”.

With the combined, batch-corrected transcriptomic data, we performed DESeq2 and identified fewer differentially expressed genes (DEGs) between high and low efficiency differentiations across cell lines (**Figure 2F, Figure S2E**) than in WTC11 hiPSC differentiation alone (**Figure 1**). These conserved differentiation step-specific DEGs were input into clusterProfiler to identify key pathways differentially active between high and low efficiency differentiations at each differentiation stage (**Figure 2G**)^39^. From this analysis we note a few significant terms: at the +IWP2 step, the high efficiency group was enriched for “cAMP” and “calcium signaling”; at the −IWP2 step, the low efficiency group was enriched for “cAMP”, “p53”, and “MAPK signaling” in addition to the cardiac-specific term “hypertrophic cardiomyopathy”; and at the +INS step, the high efficiency group was enriched for “calcium”, “cGMP-PGK”, and “insulin signaling” pathways and the low-efficiency group was enriched in “TGF-β signaling”.

Reassessing the combined, batch-corrected transcriptomic data in conjunction with the omics data from the original WTC11 hiPSC dataset (**Figure 1**), we investigated the number of features from each -ome exhibiting differential abundance between high and low efficiency batches at each differentiation step (**Figure 2H**). Although a majority of features were distinct at each step (30, 830, 792, and 1234 differential features at the +CHIR, +IWP2, −IWP2, and +INS steps, respectively), several features were shared among various steps. Four features were differentially abundant at all steps; these included two noncoding RNAs (LOC107987258 and LOC107985147) and the genes *PCA3* and *ZNF229* with the first 3 enriched in low efficiency batches and *ZNF229* enriched in high-efficiency batches (**Figure S2F**). There were more features (34 features) that were differentially abundant at all points except the +CHIR differentiation step because there were few total differential features (65 features) present at the +CHIR differentiation step (**Figure 2H**, **Figure S2G**). Among these 34 features differentially abundant at all but the +CHIR differentiation step, 11 genes were conserved as enriched in the high efficiency group and the STRING network assembled from these identified *ADCY5* and *PPP1R1B* (genes involved in cAMP signaling) as central nodes (**Figure S2H**). Interestingly, the +IWP2-specific features were more heavily driven by proteins than most other groups. Additionally, metabolites appeared to describe earlier differences between the groups, representing a much greater proportion of the differentially abundant features at the +CHIR and +IWP2 differentiation steps than later steps, as well as being among the most differentially expressed at all timepoints (**Figure 2H-I, Figure S2I**).

To capture pathway-level differences between high and low efficiency differentiations, we performed comprehensive gene set enrichment and overrepresentation analyses across KEGG, GO, and Reactome pathways, alongside upstream regulator prediction (**Figure 2J**)^40^. Notably, most of the top differential RNAs and metabolites were ncRNAs and lipid species respectively, with both predominantly enriched in low efficiency differentiations. Furthermore, several pathways switched in their enrichment across time (for example, “NF-KB” was enriched in high efficiency differentiations at the +IWP2 step but at the −IWP2 step it was enriched in the low efficiency group), highlighting the temporal and contextual dependence of these pathways in proper fate acquisition. We were particularly interested in the differentially expressed pathways and predicted transcriptional regulators at key differentiation steps as they provided crucial insight into testable hypotheses aimed at improving the differentiation efficiency of sub-optimal hPSC-CM differentiation.

### Differential responses to differentiation induction drive early multi-omic divergence in hPSC-CM differentiation efficiency

Motivated by the significant early divergence in response to CHIR-based CM differentiation induction, which was apparent by the +IWP2 stage of the differentiation, we sought to leverage the multi-omics dataset to more deeply investigate the temporal dynamics in divergent cell populations from the +CHIR to the +IWP2 differentiation step. To do this, we first performed differential analysis on features changing between these two steps within the high and low efficiency groups independently and compared these dynamic profiles to each other (**Figure 3A**, comparisons 1 and 2). The differential feature table and upset plot (**Figure 3B**) as well as the parity plot (**Figure S3A**) highlight significant overlaps in the trajectories of high and low efficiency CM differentiations as they transitioned from the +CHIR to the +IWP2 differentiation step. Notably, a majority of features changing from the +CHIR step to the +IWP2 step were shared in both high and low efficiency batches (4995 features). Conversely, there were fewer, though still many, features unique to high (2212 features) or low (1908 features) efficiency differentiation groups during the temporal transition from the +CHIR to the +IWP2 differentiation step.

**Figure 3.**
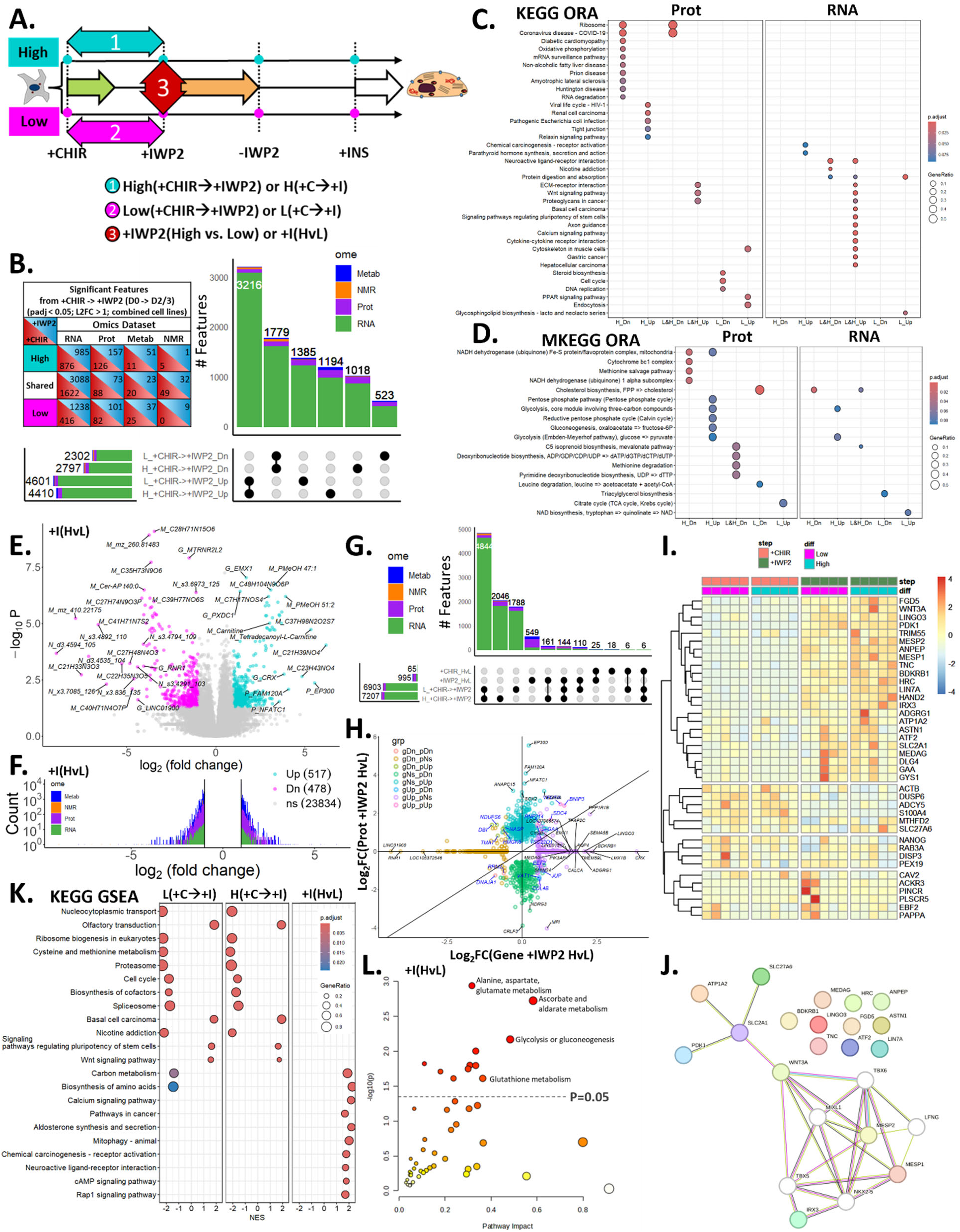
Differential responses to differentiation induction drive early multi-omic divergence in hPSC-CM differentiation efficiency. **A)** Schematic of the hPSC-CM differentiation steps and three comparisons of interest for the +CHIR to +IWP2 transition. **B)** Differential features from the +CHIR to +IWP2 differentiation step unique to high or low efficiency differentiations or shared among high and low efficiency differentiations. The number of differentially expressed features for each comparison are represented in a table (left) and an UpSet plot (right) where they are further separated by direction of change where Up corresponds to enriched and Dn (Down) corresponds to decreased for the temporal comparison as indicated. **C-D)** Overrepresentation analysis for differentially expressed proteomic and transcriptomic features for the **C)** KEGG and **D)** MKEGG databases. Comparisons are denoted as H (high efficiency), L (low efficiency), Up (enriched), and Dn (decreased) on the x-axis. Comparisons shown are enriched during the +CHIR to +IWP2 transition in either high, low, or both efficiency groups as indicated. **E)** Volcano plot of the differential features detected at the +IWP2 step between high (turquoise) and low (magenta) efficiency groups. Prefixes for each -omic feature are G = gene, P = protein, M = intracellular metabolite, N = extracellular metabolite. **F)** Histogram for the number of differentially abundant -omic features with the corresponding log2 fold change between high and low efficiency differentiation groups at the +IWP2 step. **G)** UpSet plot of differential features for the three comparisons in Panel A as well as for the comparison between high and low differentiation efficiency groups at the +CHIR differentiation step. **H)** Parity plot of the log2 fold change for genes (x-axis) and proteins (y-axis) differentially expressed between high and low efficiency groups at the +IWP2 differentiation step. **I)** Heatmap of RT-qPCR data of 39 of the top differentially expressed genes at the +IWP2 differentiation step from transcriptomics assessed in 10 additional differentiation batches from WTC11 hiPSCs (5 each for high and low efficiency differentiation groups; 2^-ΔCT). **J)** STRING network of genes conserved as enriched in the high efficiency group at the +IWP2 differentiation step. **K)** Gene set enrichment analysis for differentially expressed transcriptomic features for the KEGG database. Comparisons are shown for temporal or high (H) versus low (L) efficiency comparisons for the +IWP2 differentiation step. **L)** Joint Pathway Analysis of metabolic pathways combining differential proteins and metabolites at the +IWP2 differentiation step. ΔCT = cycle threshold of gene minus cycle threshold of the geomean of 3 reference genes (*ZNF384*, *EDF1*, *DDB1*).

To functionally investigate both the shared and unique dynamics through this differentiation transition, we performed ORA for differential proteins and transcripts using the KEGG and Reactome pathway databases (**Figure 3C, Figure S3B**). We found 3 enriched KEGG pathways, including “ECM-receptor interaction”, “Wnt signaling pathway”, and “Proteoglycans in cancer” conserved among both high and low differentiation efficiency groups (**Figure 3C**). Wnt and ECM signaling are both known to play important roles in hPSC-CM fate commitment, and these data suggest that low efficiency batches do not arise simply from an absence of Wnt signaling agonism at the onset of the differentiation or the failure to remodel the ECM during the epithelial-mesenchymal transition^41,42^. Shared among high and low efficiency differentiations in both -omes we found 5 enriched Reactome pathways, including terms such as “Cardiogenesis”, “Gastrulation”, and “Integrin cell surface interactions” (**Figure S3B**).

For unique temporal dynamics, the proteome pathway results indicated increased “relaxin” and “hypoxia signaling” while the transcriptome pointed toward increased “fibroblast growth factor signaling” in the high efficiency differentiation group compared to the low efficiency group (**Figure 3C, Figure S3B**). Moreover, the proteome also suggested potential “cell cycle” differences and differential energy metabolic utilization emerging between the high and low efficiency differentiation groups from the +CHIR to the +IWP2 differentiation step (**Figure 3C, Figure S3B**). Notably, “oxidative phosphorylation” decreased in the high efficiency group and “PPAR signaling” increased in the low efficiency group (**Figure 3C**). To look deeper at these potential metabolic differences, we repeated the ORA analysis for the MKEGG database and found that both -omes suggested a shift towards dependence on “glycolysis” for the high efficiency group and priming for more “TCA cycle” utilization by the low efficiency group (**Figure 3D**).

To further investigate differences between high and low efficiency differentiation groups, we compared differentially expressed features present at the +IWP2 differentiation step (**Figure 3A**, comparison 3). Notably, there were 995 differentially expressed features between high and low efficiency groups at the +IWP2 differentiation step whereas there were only 65 differentially expressed features at the +CHIR differentiation step before the onset of differentiation (**Figure 3E-G, Figure S3C**). Interestingly, although a majority of differentially abundant features were proteins, those exhibiting the greatest fold changes between high and low efficiency groups were predominantly metabolites, further demonstrating the metabolic divergence between the groups as noted in pathway analysis (**Figure 3F-G, Figure S3C**). Because a majority of the differential features between high and low efficiency differentiation groups at the +IWP2 differentiation step were proteins, we constructed a parity plot comparing the differential expression of genes and proteins between the high and low efficiency groups (**Figure 3H**). Surprisingly, few gene-protein pairs changed expression across both -omes with most features present on the axes indicating enrichment in one -ome but not the other. In fact, only 17 shared features were differentially expressed in both -omes, with only 8 of these changing in the same direction in both the transcriptome and proteome (up in high: *VEGFA, BNIP3, RNF214, TSC22D1, GAA, SDC4*; up in low: *RRM2, DNAJA1*).

To assess the reproducibility of top DEGs at the +CHIR and +IWP2 differentiation steps, we performed RT-qPCR on a panel of 39 genes to assess gene expression on a separate set of 10 independent differentiations with 5 each for high and low efficiency differentiation groups (**Figure 3I**, **Figure S3D**). These 10 differentiations exhibited normal profiles of canonical CM differentiation marker genes with some differences between high and low efficiency differentiation groups appearing at the +INS differentiation step (**Figure S3E**). Notably, the RT-qPCR data exhibited similar trends for a majority of the top DEGs assessed (**Figure 3I**). A STRING network constructed with conserved marker genes at the +IWP2 differentiation step showed some contribution from a core cardiac developmental and related Wnt-signaling subnetwork and several independent genes which may play other roles in batch divergence (**Figure 3J**).

Combining the temporal transitions within each efficiency group and the resulting differential state between high and low efficiency groups at the +IWP2 differentiation step, we performed GSEA using the KEGG database for the three comparisons of interest (**Figure 3A,K**). Again, while a majority of the pathway changes from the +CHIR to the +IWP2 differentiation step were shared among groups, we identified two KEGG gene sets uniquely depleted in the low efficiency group, “carbon metabolism” and “biosynthesis of amino acids”. Moreover, we observed enrichment of several gene sets in the high efficiency group at the +IWP2 stage, including “carbon metabolism” and “biosynthesis of amino acids” as well as signaling pathways for “calcium”, “cAMP”, and “Rap1”. These findings highlight the utility of studying both high and low efficiency CM differentiations to understand differences between the two differentiation outcomes that can lead to batch failure. To further investigate the metabolic shifts inferred by transcriptomic and proteomic pathway analyses, we performed joint pathway analysis using MetaboAnalyst to functionally combine the proteomic and metabolomic data^43^. This analysis corroborated our previous metabolic findings, with “glycolysis”, as well as pathways involving “amino acid” and “redox metabolism”, being enriched in the high efficiency differentiation group in comparison to the low efficiency group at the +IWP2 differentiation step (**Figure 3L**).

### Multi-omic divergence between high and low efficiency hPSC-CM differentiations persists post-Wnt inhibition

After discovering major multi-omic differences between high and low efficiency hPSC-CM differentiations in response to CHIR-based differentiation induction (by the +IWP2 step), we examined whether batch divergence persisted after the completion of Wnt inhibition. To do this, we first compared how high and low efficiency differentiations individually progressed from the +IWP2 to −IWP2 differentiation step as well as how high and low efficiency groups differed at the −IWP2 differentiation step (**Figure 4A**, comparisons 1-3). Interestingly, although the high and low efficiency differentiations diverged significantly by the +IWP2 stage, they maintained substantial conservation of temporal dynamics during the +IWP2 to −IWP2 transition (**Figure 4B, Figure S4A-B**). This was clearly evident by the high degree of similarity in the parity plot for multi-omic features changing through these differentiation steps between high and low efficiency groups (**Figure S4A**, similar features along diagonal line). Notably, there were 2166 features that changed in both high and low efficiency groups during the +IWP2 to −IWP2 transition (**Figure 4B**). To examine these shared temporal dynamics, we performed GSEA and ORA representation analyses for transcriptomic and proteomic features (**Figure 4C, Figure S4C-J**). Notably, for KEGG GSEA, high and low efficiency differentiations shared enrichment for pathways such as “Cytoskeleton in muscle cells” and “TGF-beta signaling pathway” during the +IWP2 to −IWP2 transition, which indicated shared developmental signaling as well as the enrichment of structural genes related to muscle cell development (**Figure 4C**). For additional pathway analyses, high and low efficiency differentiations shared enrichment for pathways related to “cardiac muscle contraction/development”, “PI3K-Akt signaling”, “regulation of Wnt signaling”, and “ECM organization” (**Figures S4C-J**). Moreover, both efficiency groups shared depletion for pathways related to “Activin signaling”, “apoptotic signaling”, and “gastrulation” (**Figures S4C-J**)

**Figure 4.**
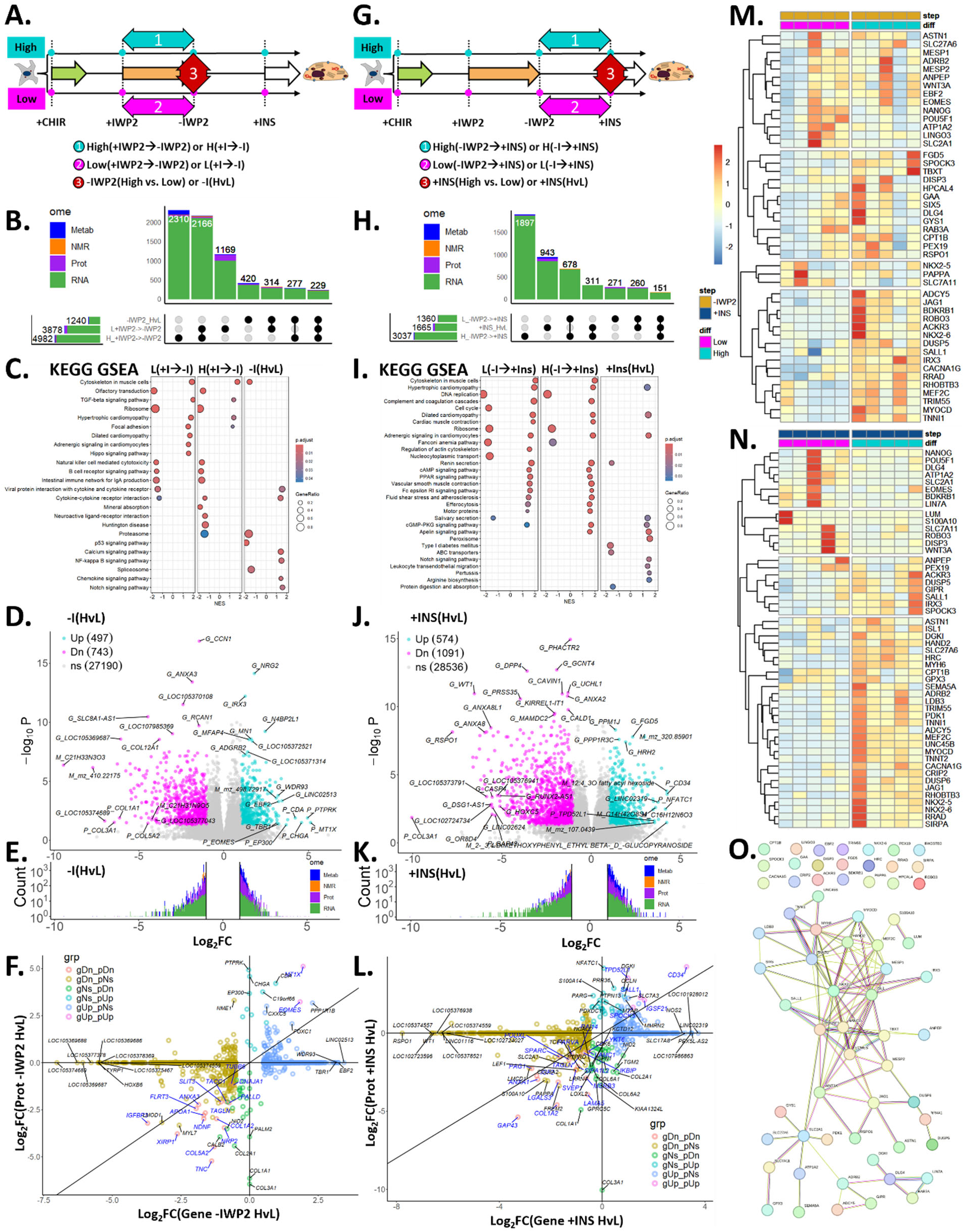
Multi-omic divergence between high and low efficiency hPSC-CM differentiations persists post-Wnt inhibition. **A)** Schematic of the hPSC-CM differentiation steps and three comparisons of interest for the +IWP2 to −IWP2 transition. **B)** UpSet plot of differential features for the three comparisons in Panel A for the +IWP2 to −IWP2 transition. **C)** Gene set enrichment analysis for differentially expressed transcriptomic features for the KEGG database. Comparisons are shown for temporal or high (H) versus low (L) differentiation efficiencies for the −IWP2 differentiation step. **D)** Volcano plot of the differential features detected at the −IWP2 step between high (turquoise) and low (magenta) efficiency groups. Prefixes for each -omic feature are G = gene, P = protein, M = intracellular metabolite, N = extracellular metabolite. **E)** Histogram for the number of differentially abundant -omic features with the corresponding log2 fold change between high and low efficiency differentiation groups at the −IWP2 step. **F)** Parity plot of the log2 fold change for genes (x-axis) and proteins (y-axis) differentially expressed between high and low efficiency groups at the −IWP2 differentiation step. **G)** Schematic of the hPSC-CM differentiation steps and three comparisons of interest for the −IWP2 to +INS transition. **H)** UpSet plot of differential features for the three comparisons in Panel G for the −IWP2 to +INS transition. **I)** Gene set enrichment analysis for differentially expressed transcriptomic features for the KEGG database. Comparisons are shown for temporal or high (H) versus low (L) differentiation efficiencies for the +INS differentiation step. **J)** Volcano plot of the differential features detected at the +INS step between high (turquoise) and low (magenta) efficiency groups. Prefixes for each -omic feature are G = gene, P = protein, M = intracellular metabolite, N = extracellular metabolite. **K)** Histogram for the number of differentially abundant -omic features with the corresponding log2 fold change between high and low efficiency differentiation groups at the +INS step. **L)** Parity plot of the log2 fold change for genes (x-axis) and proteins (y-axis) differentially expressed between high and low efficiency groups at the +INS differentiation step. **M)** Heatmap of RT-qPCR data of 46 of the top differentially expressed genes at the −IWP2 differentiation step from transcriptomics assessed in 10 additional differentiation batches (5 each for high and low efficiency differentiation groups; 2^-ΔCT). **N)** Heatmap of RT-qPCR data of 51 of the top differentially expressed genes at the −IWP2 differentiation step from transcriptomics assessed in 10 additional differentiation batches from WTC11 hiPSCs (5 each for high and low efficiency differentiation groups; 2^-ΔCT). **O)** STRING network of genes enriched in the high efficiency group at the −IWP2 and +INS differentiation steps. ΔCT = cycle threshold of gene minus cycle threshold of the geomean of 3 reference genes (*ZNF384*, *EDF1*, *DDB1*).

Despite many shared features between high and low efficiency differentiation groups during the +IWP2 to −IWP2 transition, this temporal transition had a higher proportion of differentially changing features that were unique to the high (2310 features) and low (1169) efficiency differentiation groups (**Figure 4B, Figure S4B**). In proteomic and transcriptomic pathway analyses, these temporal differences resulted in several unique pathways enriched in high efficiency differentiation during the −IWP2 to +IWP2 transition including negative regulation of “Wnt signaling”, “Calcium signaling”, and “Transcriptional regulation of pluripotent stem cells” (**Figure S4C-J**). Interestingly, for proteomic pathway analysis, “negative regulation of TCF-dependent signaling by Wnt ligand antagonists” appeared as enriched during the Wnt inhibition differentiation step for the high efficiency differentiation group but was downregulated in the low efficiency group, which suggests that inadequate Wnt inhibition could contribute to batch divergence during this transition (**Figure S4D**). For low efficiency differentiation pathway analysis during the −IWP2 to +IWP2 transition, several GOBP terms related to “cardiac muscle tissue development’ were enriched in the proteomics data, which adds further support to the concept that low efficiency hPSC-CM differentiation may progress at faster rate (**Figure S4E**).

To further explore differences between high and low efficiency differentiation groups, we compared differentially expressed features present at the −IWP2 differentiation step (**Figure 4A**, comparison 3). Predictably, there were more differential features between high and low efficiency differentiation groups present at the −IWP2 step (1240 features) in comparison to the previous +IWP2 step (995 features) as differences expanded over time (**Figure 4B, D-E**). Surprisingly, differences at the −IWP2 differentiation step were dominated by the transcriptome as opposed to the proteome-driven differences at the +IWP2 differentiation step (**Figure 4B, E**). In contrast to the +IWP2 differentiation step, we noted more similarity in gene and protein expression differences at the −IWP2 differentiation step with 18 gene-protein pairs sharing significantly different expression between high and low efficiency groups (**Figure 4F**, similar features along diagonal line). This included high efficiency differentiation enrichment in a metal ion binding protein MT1X and a mesodermal development marker EOMES and low efficiency differentiation enrichment in an insulin growth factor binding protein IGFBP3 and an ECM protein component COL1A2. In pathway analysis, several terms were enriched in the high efficiency group in comparison to the low efficiency group at the −IWP2 stage including “calcium”, “NF-kappa B”, “Notch”, and “PI3K/ERBB4 signaling” as well as “mesoderm” and “blood vessel development” (**Figure 4C**, **Figure S4F-I**). Additionally, MKEGG metabolic pathway analysis indicated a continued enrichment of “glycolysis” in the high efficiency group in comparison to the low efficiency group, which was in line with results at the +IWP2 step (**Figure S4J**).

To continue our investigation of hPSC-CM batch divergence during post-Wnt inhibition differentiation stages, we compared the −IWP2 to +INS temporal transition for high and low efficiency differentiations individually. Additionally, we directly compared the high and low efficiency groups at the +INS differentiation step (**Figure 4G**, comparisons 1-3). Unlike previous differentiation transitions, we found far fewer features (678 features) shared between high and low efficiency groups during the −IWP2 to +INS transition (**Figure 4H, Figure S5A-B**). Notably, 408 features were shared as upregulated, while only 276 features were shared as downregulated during this transition among high and low efficiency groups (**Figure S5B**). Because a majority of these shared features were differentially expressed transcripts, we utilized pathway analysis to understand potential shared functions. From these analyses, we found that over the −IWP2 to +INS transition, high and low efficiency groups shared enrichment in terms such as “cytoskeleton in muscle cells” (continued from prior differentiation transition), “cardiac muscle contraction”, and “ECM organization” as well as “cAMP”, “cGMP”, and “PPAR signaling” (**Figure 4I**, **Figure S5C-J**). These data demonstrate that even in the context of unique differentiation outcomes, there are many shared cellular functions and signaling pathways as cardiac progenitor cells give rise numerous cardiac cell types.

While similarities existed between high and low efficiency differentiations during the −IWP2 to +INS transition, there were far more differences between high and low efficiency groups during this transition than prior differentiation transitions. For instance, there were 1897 features unique to the high efficiency group and 271 features unique to the low efficiency group during the −IWP2 to +INS transition while only 678 features were shared between groups (**Figure 4H**). Because of the clear temporal divergence between high and low efficiency groups from the −IWP2 to +INS step, we again employed pathway analysis to annotate functions to differentially changing proteins and genes. From this temporal analysis, “apelin signaling” and “cholesterol metabolism” were significantly enriched specifically in the high efficiency group whereas “glutathione metabolism”, “mesenchymal cell differentiation”, and “Wnt signaling” were depleted (**Figure 4I**, **Figure S5C-I**). Several pathways were also unique to the low efficiency −IWP2 to +INS temporal transition including an enrichment in “cardiac muscle contraction” and “cardiac morphogenesis” at the protein level and a depletion in “cell cycle”, “cholesterol biosynthesis”, and “lipid metabolism” (**Figure 4I, Figure S5C-J**). These data again support an earlier acquisition of the cardiomyocyte-related proteins and phenotypes as a potential indicator of batch failure.

To more fully harness the utility of characterizing high and low efficiency hPSC-CM differentiation batches, we directly compared these groups at the +INS differentiation step (**Figure 4G**, comparison 3). Unsurprisingly, the number of differential features between high and low efficiency groups present at the +INS step (1665 features) continued to expand in comparison to the previous −IWP2 (1240 features) and +IWP2 (995 features) differentiation steps (**Figure 4H, J-K**). Similar to the −IWP2 step, differences at the +INS differentiation step were dominated by the transcriptome (**Figure 4H, K**). Moreover, we noted slightly increased similarity in gene and protein expression differences at the +INS differentiation step in comparison to the −IWP2 step with 23 gene-protein pairs sharing significantly different expression between high and low efficiency differentiation groups (**Figure 4L**, similar features along diagonal line). This included high efficiency differentiation enrichment in endothelial cell marker CD34 and cardiac precursor marker SALL1 and low efficiency differentiation enrichment in ECM protein components COL1A2 (continued from prior step) and LAMA5. For pathway analysis, several terms were enriched in the high efficiency group in comparison to the low efficiency group at the +INS stage including “negative regulation of Notch signaling”, “carbohydrate metabolism”, “glycolytic process”, and “glycolysis” (**Figure 4I**, **Figure S5F-J)**. Additionally, several terms were downregulated in the high efficiency group in comparison to the low efficiency group at the +INS stage including “TGF-beta signaling” and “ECM organization” (**Figure 4I**, **Figure S5F-J**). Together, these pathway results indicate unique signaling to high and low efficiency differentiation batches at the +INS step. Furthermore, these findings demonstrate a continued metabolic separation in line with findings from prior differentiation steps where high efficiency batches were enriched for glycolysis-related terms.

To assess the reproducibility of top DEGs at the −IWP2 and +INS differentiation steps, we performed RT-qPCR on a panel of 46 and 51 genes respectively for a separate set of 10 independent differentiations with 5 high and 5 low efficiency differentiations (**Figure 4M-N**). For a majority of the top DEGs identified from transcriptomics, the RT-qPCR data exhibited similar trends. Notably, there was significant overlap between the top DEGs for the −IWP2 and +INS differentiation steps. Because of this, we created a single STRING network to assess how the top DEGs may potentially interact (**Figure 4O**). This network demonstrated significant interactions between known key cardiac developmental regulators such as *NKX2-5*, *ISL1*, and *MESP1* as well as several subnetworks related to less characterized regulators such as *SLC2A1* and *JAG1*. Interestingly, there were also many network-independent entities including *RHOBTB3*, *HRC*, and *CPT1B* with unknown interactions to key cardiac developmental regulators.

### Metabolic differences distinguish high and low efficiency progenitors during early hPSC-CM differentiation

After observing significant metabolic differences between high and low efficiency hPSC-CM progenitors in the transcriptomic, proteomic, and intracellular metabolomic data, we set out to further characterize metabolism during early differentiation steps. To do this, we first used MetaboAnalyst to map 117 out of the 1522 intracellular metabolites to the Human Metabolome Database (HMDB) for PCA, differential abundance, and metabolite set enrichment analysis at the +CHIR, +IWP2, −IWP2, and +INS differentiation steps. At the +CHIR and −IWP2 differentiation steps, there were only 0 and 1 differentially abundant compounds between high and low efficiency differentiation groups respectively. Moreover, there were no significantly different pathways by metabolite set enrichment analysis (MSEA) using the SMPDB and RaMP-DB pathway databases.

For intracellular metabolomics at the +IWP2 differentiation step, PCA analysis demonstrated significant separation (PERMANOVA p-value < 0.05) between high and low efficiency groups (**Figure 5A**). Additionally, there were 2 compounds significantly enriched in the low efficiency differentiation group and 9 compounds significantly enriched in the high efficiency group (Log_2_FC > 1, FDR adjusted p-value < 0.05, **Figure 5B**). Using quantitative enrichment analysis, we identified 50 SMPDB pathways and 103 RaMP-DB pathways with significant differences between high and low efficiency differentiation groups at the +IWP2 differentiation step (FDR adjusted p-value < 0.05, **Figure 5C**, **Figure S6A**). In the top 25 pathways of SMPDB and RaMP-DB MSEA analysis, there were numerous pathway changes related to “Fatty acid Metabolism”, which were predominantly driven by changes in L-carnitine and L-acetylcarnitine species.

**Figure 5.**
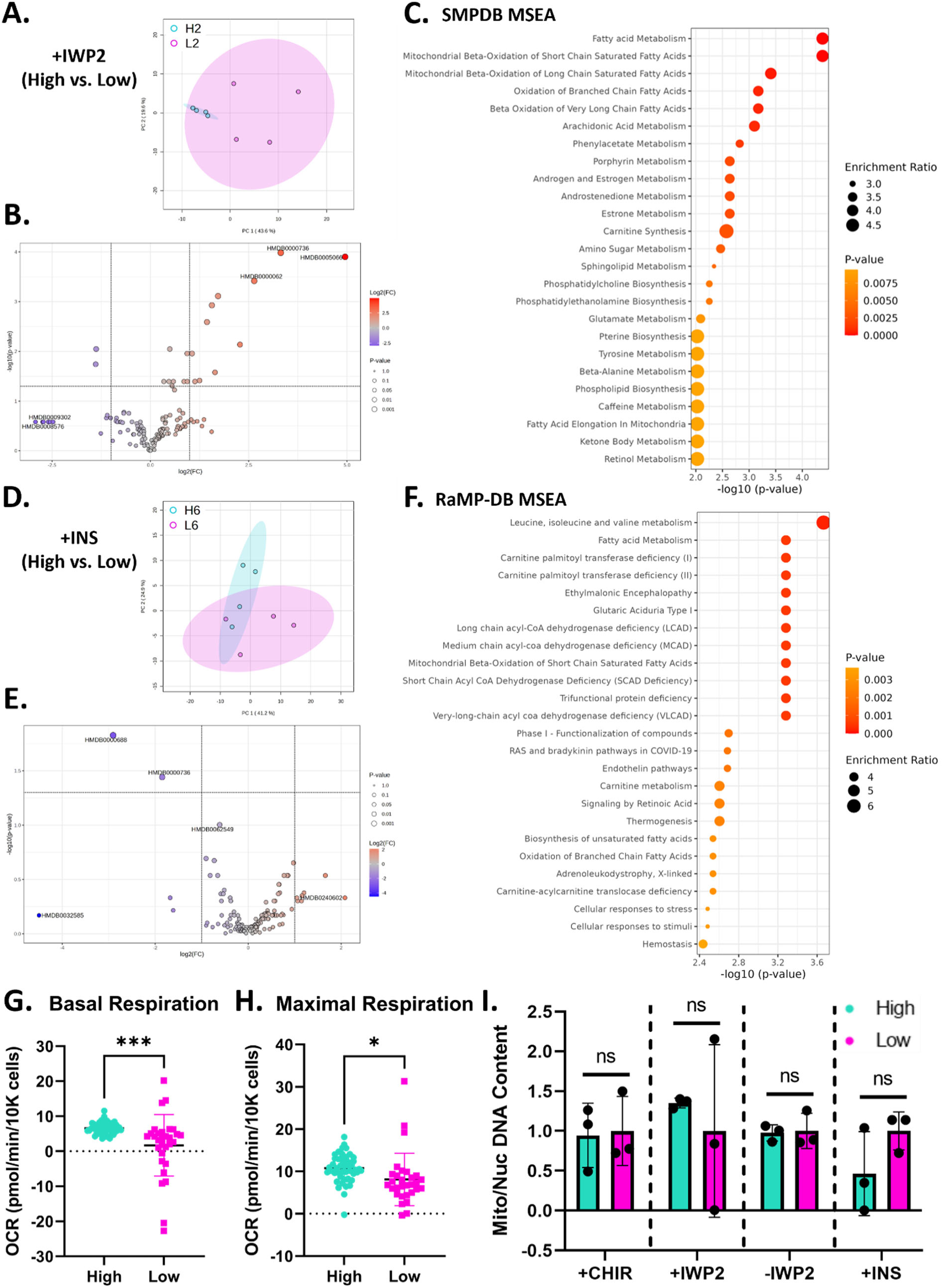
Metabolic differences distinguish high and low efficiency progenitors during early hPSC-CM differentiation stages. **A)** PCA of intracellular metabolomes at the +IWP2 step (D2) for high (turquoise, H2) and low (magenta, L2) differentiation efficiencies (95% CI, n = 4 well replicates per group). **B)** Volcano plot of differentially abundant intracellular metabolites enriched at the +IWP2 step, enriched in high (red) or low (blue) efficiency groups (Log2FC > 1, FDR adjusted p-value < 0.05). **C)** Metabolite set enrichment analysis (MSEA) of intracellular metabolites using the SMPDB database displaying the top 25 pathways enriched in high efficiency progenitors at the +IWP2 differentiation step (all with FDR adjusted p-value < 0.05, raw p-values shown). **D)** PCA of intracellular metabolomes at the +INS step (D6) for high (turquoise, H6) and low (magenta, L6) differentiation efficiencies (95% CI, n = 4 well replicates per group). **E)** Volcano plot of differentially abundant intracellular metabolites enriched at the +INS step, enriched in high (red) or low (blue) efficiency groups (Log2FC > 1, FDR adjusted p-value < 0.05). **F)** MSEA of intracellular metabolites using the RaMP-DB database displaying the top 25 pathways enriched in high efficiency progenitors at the +INS differentiation step (all with FDR adjusted p-value < 0.05, raw p-values shown). **G-H)** MitoStress test at +INS differentiation step (D6) for cryopreserved cardiac progenitor cells from high (turquoise, n = 46 well replicates) and low (magenta, n = 30 well replicates) efficiency hPSC-CM differentiations from WTC11 hiPSCs. Unpaired t-tests compare **G)** basal and **H)** maximal respiration measured by Seahorse OCR. **I)** Mitochondrial-to-nuclear DNA content (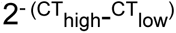) normalized to the step-specific low efficiency cardiomyocyte differentiation average for high (turquoise) and low (magenta) efficiency cardiomyocyte differentiations across +CHIR (D0), +IWP2 (D2), −IWP2 (D4), and +INS (D6) steps in WTC11 hiPSCs (n=3 differentiation replicates). High efficiency differentiations (>63% cTnT+ cells) and low efficiency differentiations (<32% cTnT+ cells) shown. P-values from unpaired t-tests. All data are represented as mean ± SD.

At the +INS differentiation step, PCA analysis of intracellular metabolites demonstrated significant separation (PERMANOVA p-value < 0.05) between high and low efficiency groups (**Figure 5D**). Additionally, there were 2 compounds significantly enriched in the high efficiency group (Log2FC > 1, FDR adjusted p-value < 0.05, **Figure 5E**). Using quantitative enrichment analysis, we identified 1705 RaMP-DB pathways with significant differences between high and low efficiency differentiation groups at the +INS differentiation step (FDR adjusted p-value < 0.05, **Figure 5F**). In the top 25 pathways of RaMP-DB MSEA analysis, there were again numerous pathways related to “Fatty acid Metabolism”. To functionally interrogate metabolic differences between high and low efficiency hPSC-CM progenitors at the +INS differentiation step, we performed a MitoStress test. Using this approach, we assessed differences in respiration between high and low efficiency differentiations from oxygen consumption rate (OCR) measurements (**Figure S6B**). From this assessment, high efficiency hPSC-CM progenitors exhibited significantly increased basal, maximal, and ATP-linked respiration compared to low efficiency hPSC-CM progenitors (**Figure 5G-H, Figure S6C**). Conversely, low efficiency hPSC-CM progenitors exhibited significantly increased non-mitochondrial oxygen consumption (**Figure S6D**). Moreover, increases in respiration parameters in high efficiency hPSC-CM progenitors at the +INS differentiation step were not due to significant differences in mitochondrial/nuclear DNA content at any of the early differentiation steps (**Figure 5I**).

To further examine metabolic differences between high and low efficiency hPSC-CM progenitors, we performed Nuclear Magnetic Resonance (NMR) extracellular metabolomics on spent media from high and low efficiency differentiations at early differentiation steps. From 1D and 2D NMR spectra, we identified and quantified 25 metabolites and performed PCA, differential abundance analysis, and metabolite set enrichment analysis at the +CHIR, +IWP2, −IWP2, and +INS differentiation steps. At the +CHIR step, there were no significantly different extracellular metabolites or pathways by metabolite set enrichment analysis (MSEA) using the SMPDB and RaMP-DB pathway databases. At the +IWP2 differentiation step, there was significant group separation on PCA (PERMANOVA p-value < 0.05), and glucose was significantly enriched in the low efficiency differentiation group (Log_2_FC > 1, FDR adjusted p-value < 0.05, **Figure S6E-F**). Enrichment of glucose in the media of the low efficiency differentiation group reflects reduced glucose utilization, which is supported by the SMPDB MSEA analysis demonstrating enrichment of “Glycolysis” in the high efficiency differentiation group (**Figure S6G**). At the −IWP2 differentiation step, glucose enrichment in the low efficiency group was conserved and a number of similar SMPDB pathways including “Glycolysis” and “Gluconeogenesis” were significantly enriched in the high efficiency differentiation group (**Figure S6F)**. At the +INS differentiation step, there was again significant group separation on PCA (PERMANOVA p-value < 0.05, **Figure S6H**). SMPDB MSEA analysis again demonstrated enrichment of “Gluconeogenesis” in the high efficiency differentiation group, which was driven by a significant increase in glucose in the low efficiency differentiation media and a significant increase in lactate in the high efficiency differentiation media (FDR adjusted p-value < 0.05 but Log2FC < 1, **Figure S6I**). Together, these NMR extracellular metabolomic data indicated an enrichment in glycolysis in high efficiency differentiation progenitors, which was consistent with earlier transcriptomic and proteomic findings. In addition, the intracellular metabolomic and oxygen consumption data demonstrated an increase in oxidative respiration in high efficiency differentiation progenitors. In total, our metabolic investigations suggested a combination of increased glycolytic and oxidative metabolism in high efficiency hPSC-CM differentiation progenitors in comparison to low efficiency hPSC-CM differentiation progenitors.

### Machine learning models using early conserved DEGs quantitatively predict hPSC-CM differentiation purity

After identifying differentially expressed genes in high and low efficiency hPSC-CM differentiations at early differentiation stages, we asked whether expression of these genes could be used to develop models that quantitatively predict terminal cardiomyocyte purity (% cTnT+ cells) from progenitor cell populations. Toward this goal, we identified an initial list of 54 genes from transcriptomics that were validated with RT-qPCR as predictive in distinguishing high and low efficiency hPSC-CM differentiation batches at one or more differentiation steps (**Figure 6A**, **Figure S7A-B**). Of these 54 genes, 11 were also detected within the proteomics dataset, all of which observed qualitatively similar enrichment trends as our transcript-level findings (**Figure S7C**). We then performed RT-qPCR to quantify expression of these 54 genes of interest across 3 hPSC lines for all 4 differentiation steps (+CHIR, +IWP2, −IWP2, +INS) in 22 additional hPSC-CM differentiation batches with terminal CM purities ranging from 5-81% cTnT+ cells. Notably, 31 of these genes demonstrated quantitative predictive ability at one or more differentiation steps in RT-qPCR analysis (**Figure S7B**). To further validate the predictive power of these 31 genes, we queried a publicly-available time course transcriptomic dataset comprising 19 hiPSC-CM batches with a wide range of CM purity outcomes from 1.1-83.9% cTnT+ cells.^20^ We extracted the relevant timepoints that mapped onto our samples including Days 0, 3, 5, and 7 corresponding to the hPSC-CM differentiation protocol for GiWi hPSC lines. After confirming similar differentiation trajectories based on canonical markers, we found that a majority of our 31 putative predictor genes exhibited similar enrichment trends (**Figure S7D**). A STRING network of these 31 genes with CM-predictive capacity demonstrated a core of known cardiac developmental regulators and interaction of this core with various glucose and ion modulators in addition to several network-independent entities (**Figure 6B**). Interestingly, 3 of the 31 markers including *IRX3, JAG1,* and *HRC* exhibited predictive power at 3 differentiation steps (+IWP2, - IWP2, and +INS) in RT-qPCR and publicly available transcriptomic dataset analysis (**Figure 6C-D**, **Figure S7E-F**).

**Figure 6.**
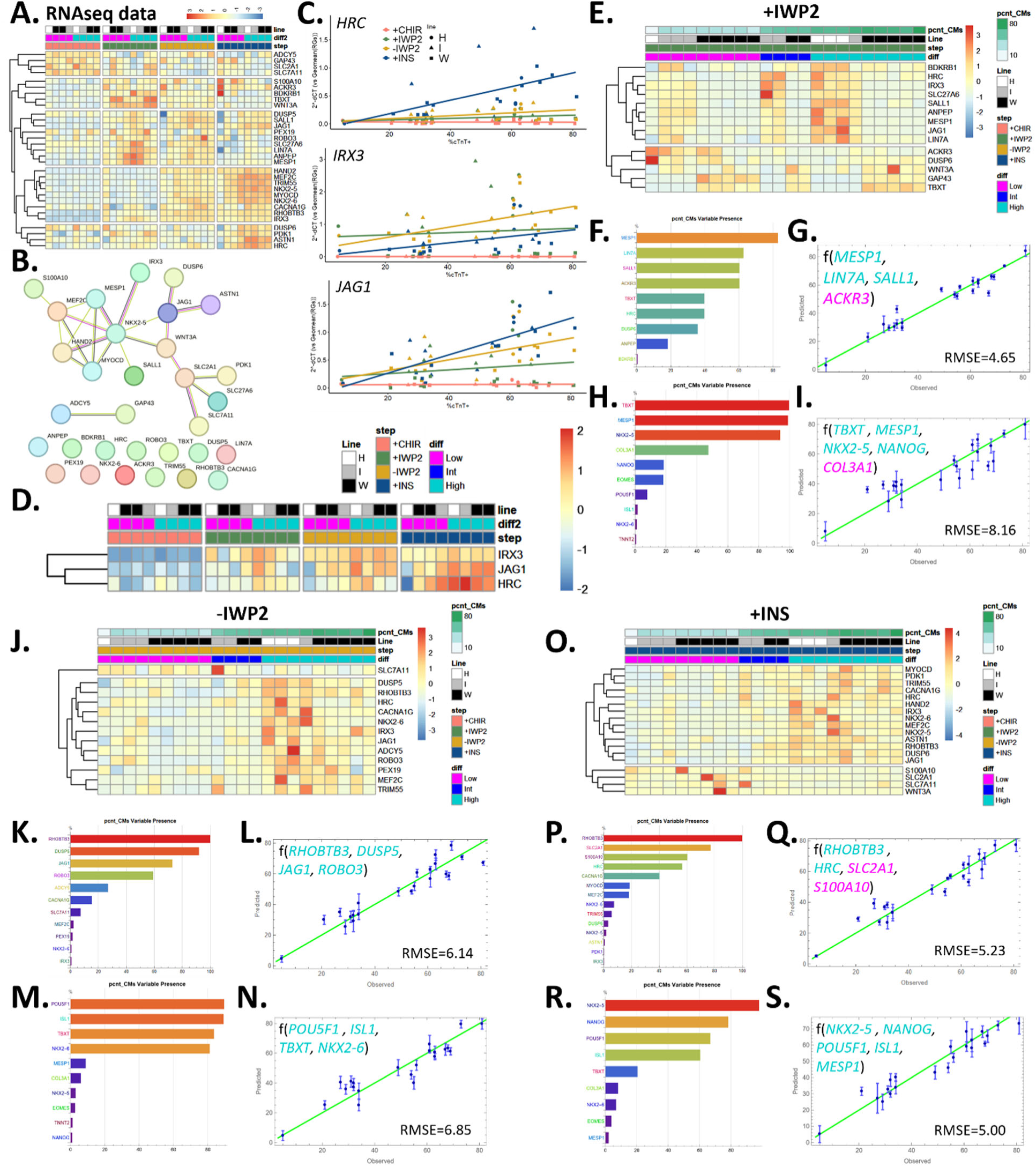
Machine learning models using early conserved DEGs quantitatively predict hPSC-CM differentiation purity. **A)** Heatmap of transcriptomic data (TPM) of 31 predictive genes across differentiation steps in 3 hPSC lines. **B)** STRING network of the 31 marker genes distinguishing high and low efficiency hPSC-CM differentiations. **C)** Regression plots of *HRC*, *IRX3*, and *JAG1* expression (qPCR 2^-ΔCT vs. %cTnT⁺) at each step in 3 hPSC lines. **D)** Heatmap (TPM) of conserved predictor genes (*HRC*, *IRX3*, *JAG1*) across differentiation steps for 3 hPSC lines. **E-I**) Modeling results for +IWP2 differentiation step. **E)** Heatmap (RT-qPCR) of 14 step-specific predictor genes. **F)** Variable presence in evolved models after one round of predictive modeling. **G)** Nonlinear prediction plot for the top 4 predictive marker genes after several rounds of model evolution. **H)** Variable presence in evolved models after one round of predictive modeling for 10 canonical cardiomyocyte differentiation markers. **I)** Nonlinear prediction plot for the top 5 canonical marker genes after one round of model evolution. **J-N)** Modeling results for −IWP2 differentiation step. **J)** Heatmap (RT-qPCR) of 13 step-specific predictor genes. **K)** Variable presence in evolved models after one round of predictive modeling. **L)** Nonlinear prediction plot for the top 4 predictive marker genes after several rounds of model evolution. **M)** Variable presence in evolved models after one round of predictive modeling for 10 canonical cardiomyocyte differentiation markers. **N)** Nonlinear prediction plot for the top 4 canonical marker genes after one round of model evolution. **O-S)** Modeling results for +INS differentiation step. **O)** Heatmap (RT-qPCR) of 18 step-specific predictor genes. **P)** Variable presence in evolved models after one round of predictive modeling. **Q)** Nonlinear prediction plot for the top 4 predictive marker genes after several rounds of model evolution. **R)** Variable presence in evolved models after one round of predictive modeling for 9 canonical cardiomyocyte differentiation markers (excluding *TNNT2*). **S)** Nonlinear prediction plot for the top 5 canonical marker genes after one round of model evolution. For nonlinear prediction plots, f(*genes*) denotes predictive functions with turquoise genes enriched in high efficiency differentiations and magenta genes enriched in low efficiency differentiations. TPM = transcripts per million. ΔCT = cycle threshold of gene minus cycle threshold of the geomean of 3 reference genes (*ZNF384*, *EDF1*, *DDB1*). pcnt_CMs = percentage of CMs (cTnT flow cytometry). RMSE = root mean squared error. Line = cell line (H = H9, I = IMR90-4, W = WTC11).

To develop differentiation step-specific predictive models for terminal cardiomyocyte purity, we prioritized top differentiation step-specific markers at the +IWP2 (14 genes), −IWP2 (13 genes), and +INS (18 genes) steps from transcriptomics (**Figure S6G-I**). We first attempted linear modeling using partial least squares regression (PLSR) with the top 54 predictor genes (including several canonical genes). Although these PLSR models were able to predict terminal purity for some differentiation batches, the discrepancy between the measured and predicted terminal cardiomyocyte purity was higher than 30% in some cases across all 3 differentiation steps (**Figure S8A-C**). Additionally, we assessed the predictive ability of 10 canonical hPSC-CM differentiation markers across all 3 differentiation steps (**Figure S8D-M**). None of these markers individually exhibited the ability to quantitatively predict terminal cardiomyocyte purity across multiple differentiation steps, and only a few markers such as *TNNT2*, *TBXT*, and *NKX2-5* displayed predictive power for one differentiation step. Because of the pitfalls encountered with linear modeling and individual canonical gene prediction of cardiomyocyte purity, we pivoted to utilizing nonlinear modeling methods afforded by DataModeler (Evolved Analytics) across the +IWP2, −IWP2, and +INS differentiation steps.

For each differentiation step, we generated predictive model ensembles from our top predictor genes and iterated with the highest model-driving variables. To understand how these models fared against canonical hPSC-CM differentiation markers, we compared the differentiation step-specific models to models generated with a panel of 10 canonical hPSC-CM differentiation markers (**Figure S7B**, **Figure S8D-M**). For the +IWP2 differentiation step, RT-qPCR data from the 14 top DEGs broadly showed gene expression changes that corresponded with CM purity across high, intermediate, and low efficiency groups (**Figure 6E**). After the first round of model generation, 9 driving variables were identified and used for model iteration (**Figure 6F**). Subsequently, 4 genes including *MESP1*, *LIN7A*, *SALL1*, and *ACKR3* were able to generate a predictive model with a root mean squared error (RMSE) = 4.65 (**Figure 6G**). Of the 4 genes, all were upregulated in high efficiency differentiations (turquoise) except for *ACKR3* (magenta), which was upregulated in low efficiency differentiations. The model derived only from canonical genes identified 5 genes, including *TBXT*, *MESP1*, *NKX2-5*, *NANOG*, and *COL3A1,* capable of generating a predictive model; however, the model performed markedly worse than the top predictor gene model with an RMSE = 8.16 (**Figure 6H-I**).

At the −IWP2 differentiation step, RT-qPCR data from the 13 top DEGs showed gene expression changes that correlated with CM purity; however, there appeared to be fewer differences between low and intermediate differentiation purities (**Figure 6J**). After the first round of model generation, 11 driving genes were identified and used for model iteration (**Figure 6K**). Subsequently, 4 genes that were all enriched in high purity differentiations (turquoise) including *RHOBTB3*, *DUSP5*, *JAG1*, and *ROBO3* were identified as being able to generate a predictive model with an RMSE = 6.14 (**Figure 6L**). The model derived only from canonical genes identified 4 genes, including *POU5F1*, *ISL1*, *TBXT*, and *NKX2-6,* capable of generating a predictive model with a slightly higher RMSE = 6.85 (**Figure 6M-N**).

At the +INS differentiation step, RT-qPCR data from the 18 top DEGs showed gene expression changes that corresponded with CM purity (**Figure 6O**). After the first round of model generation, 14 driving genes were identified and used for model iteration (**Figure 6P**). Subsequently, 2 genes that were enriched in high efficiency differentiations (*RHOBTB3* and *HRC*) and 2 genes that were enriched in low efficiency differentiations (*SLC2A1* and *S100A10*) were identified as being able to generate a predictive model with an RMSE = 5.23 (**Figure 6Q**). The model derived from 9 canonical genes excluding *TNNT2* identified 5 genes including *POU5F1*, *ISL1*, *MESP1*, *NKX2-5*, and *NANOG* capable of generating a predictive model with a similar, but slightly better RMSE=5.00 than the top predictor gene model (**Figure 6R-S**). However, the model derived from all 10 canonical genes including *TNNT2* performed the best and had an RMSE = 3.44 (**Figure S8N-O**). This model included *TNNT2*, which exhibited its earliest expression by the +INS differentiation step in addition to other canonical markers including *POU5F1*, *ISL1*, *NKX2-5*, and *EOMES*.

Using an RT-qPCR approach to validate top DEGs from our transcriptomic datasets, we were able to generate highly predictive nonlinear models from 4 genes at each key differentiation step. These predictive models performed better than canonical gene models at both the +IWP2 and −IWP2 differentiation steps. However, the top predictor gene models performed slightly worse than canonical gene models at the +INS differentiation step. In total, our models encompassed 11 total genes across the three key differentiation steps as *RHOBTB3* was shared as a predictive feature in the models for both the −IWP2 and +INS differentiation steps. These master 11 predictor genes underscore our previous analyses implicating Notch *(JAG1*), MAPK (*DUSP5*), calcium (*S100A10* and *HRC*), and glucose metabolism (*SLC2A1*) as potent pathways marking CM fate commitment.

### Wnt and MAPK inhibition improve hPSC-CM differentiation purity

We leveraged pathway-level insights from our analyses to investigate mechanisms driving divergence from CM differentiation and to inform strategies for improving CM differentiation outcomes. From these analyses, results in **Figure 2** and later substantiated in **Figures 4 and 6** suggested aberrant Wnt and MAPK signaling activation at the −IWP2 and +INS steps as potential mechanisms leading to low efficiency differentiations. Thus, we hypothesized that extending the duration of Wnt inhibition with or without the addition of MAPK inhibition would improve differentiation outcomes by increasing terminal CM purity (**Figure 7A**). To test this, we supplemented media at the −IWP2 and +INS steps with IWP2 (prolonging the duration of Wnt inhibition) and/or FHPI (p38 MAPK inhibitor) and assessed the terminal percentage of cTnT+ cells via flow cytometry (**Figure 7B**). These treatments significantly increased terminal CM purity by 22%, 29%, and 39% (relative % over control differentiation purity) with Wnt inhibition, MAPK inhibition, and dual treatment, respectively (**Figure 7C**). Notably, this differentiation improvement was observed across differentiations with control purities ranging from 10-60% cTnT+, demonstrating the broad applicability and robustness of these protocol modifications in enhancing CM differentiation purities. Further, we assessed terminal CM yield and found that while extension of Wnt inhibition did not affect total CM yield, MAPK inhibition and dual treatment resulted in CM yield increases of 37 and 25%, respectively (**Figure 7D-E**).

**Figure 7.**
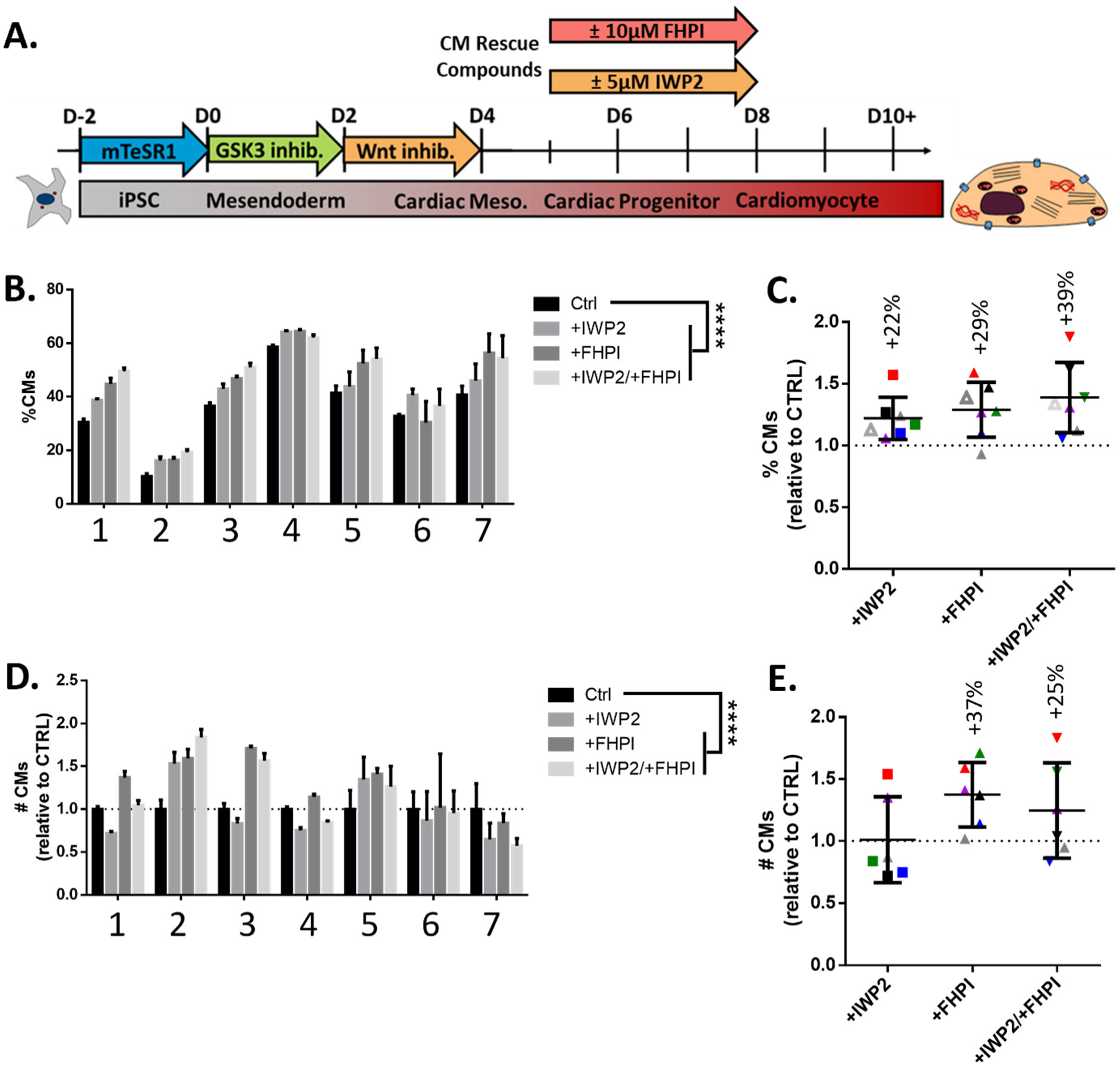
Wnt and MAPK inhibition improve hPSC-CM differentiation purity. **A)** Schematic of WTC11 hiPSC-CM differentiation showing timing, duration, and concentration of IWP2 (Wnt inhibitor), FHPI (p38 MAPK inhibitor), or dual treatment (D5–8). **B)** Terminal CM purity (% cTnT⁺ cells) from 7 independent rescue differentiations. **C)** Normalized CM purity from B); values indicate average % increase (colors denote means of 3–4 well replicates per differentiation). **D)** Terminal CM yield for the same differentiations as in (B). **E)** Normalized CM yield from D); values indicate average % increase (colors denote means of 3–4 well replicates per differentiation). P-values from one-way ANOVA with Dunnett’s post-hoc test.

Thus, our findings demonstrate that multi-omic insights from divergent differentiation trajectories can be translated into targeted process interventions. By defining both the pathways and the temporal window underlying fate divergence, we were able to intervene and suppress off-target trajectories, thereby increasing commitment to the CM lineage. More specifically, we demonstrate elevated Wnt and MAPK signaling activation as mechanisms of CM differentiation failure, which is in part rescued by Wnt and MAPK inhibition at the cardiac progenitor stage of differentiation.

## Discussion

Clinical translation of hPSC-derived therapies will require manufacturing processes that deliver consistent cell identity, purity, and therapeutic potency^44^. For, hPSC-derived cardiomyocytes (hPSC-CMs), batch-to-batch variability remains one of the many barriers to therapeutic application. Here, we used a comprehensive multi-omic strategy to define when differentiation trajectories diverge, to identify molecular and metabolic features associated with successful fate acquisition, to develop early predictive gene panels, and to implement targeted process improvements to increase CM purity and yield. Temporal comparison of high and low efficiency differentiations revealed that differentiation failure begins early, after differentiation induction with CHIR99021, a Wnt activator. Although high and low efficiency differentiations exhibited broadly similar responses to cardiogenic cues, direct contrasts uncovered distinct metabolic differences, predictive genetic features, and stage-specific signaling differences that ultimately biased lineage outcomes. More specifically, we demonstrated that high efficiency CM progenitors display functional increases in glycolytic and oxidative metabolism. Furthermore, machine learning models constructed from a panel of early gene markers were able to quantitatively predict CM purity prior to CM emergence. Lastly, stage-specific pathway analysis suggested a signaling-based mechanism for CM differentiation failure, which guided the development of a more efficient CM differentiation protocol using MAPK and sustained Wnt inhibition during the cardiac progenitor to CM transition. Together, these findings provide a framework for integrating multi-omic insights, real-time differentiation monitoring, and mechanistically informed interventions to enhance the robustness of hPSC-CM manufacturing.

Although CM differentiation protocols are highly variable, the mechanisms driving divergent differentiation outcomes remain incompletely understood^45–47^. Our multi-omic findings strengthen emerging evidence that subtle differences in hPSC responses to differentiation induction with Wnt activation dictate mesendoderm specification and downstream CM fate^46,48–51^. Multi-omic pathway analysis at the +IWP2 step demonstrated increases in FGF, calcium, MAPK, and Wnt signaling in high efficiency CM progenitors. FGF2 signaling through MEK-ERK prolongs *NANOG* expression to promote mesendoderm differentiation^52,53^, which is consistent with our observation of prolonged *NANOG* expression and an hPSC-specific gene signature in high efficiency CM differentiations despite the absence of exogenous FGF2. While Wnt activation alone can induce mesodermal differentiation in the absence of other developmental signaling gradients, correct patterning along the canonical CM developmental trajectory progressing from middle-anterior primitive streak through lateral plate mesoderm, splanchnic mesoderm, cardiac mesoderm, cardiac progenitors, and ultimately CMs is variable^14,15,52–57^. CM differentiation induction with Wnt activation is highly dependent on interactions between cell density^13,57,58^, cell cycle^47,59^, and cell death^47,60^. Higher cell densities favor CM differentiation and are associated with decreased cell cycle progression^47,61,62^. Moreover, CDK8 cell cycle inhibition improves CM differentiation reproducibility by broadening the effective CHIR99021 concentration window^59^. Cell density and death further modulate autocrine and paracrine signaling mechanisms, such as TGFβ activation and nucleotide signaling^14,57,60,63^, which are critical for robust CHIR-based CM differentiation.

Across post-Wnt inhibition differentiation stages, even less is known about CM differentiation failure modes. At the −IWP2 differentiation step, we noted increases in Notch activation and Wnt inhibition in high efficiency CM progenitors and increases in insulin growth factor (IGF) signaling in low efficiency progenitors. Increased Wnt signaling inhibition in high efficiency CM progenitors reflects a proper response to IWP2 treatment, suggesting that inadequate Wnt inhibition in low efficiency progenitors is a potential CM differentiation failure mechanism. Notch signaling activation plays a context-dependent role in hPSC-CM differentiation with evidence supporting inhibition of early mesoderm specification^64–66^ and variable effects on subsequent cardiac progenitor transitions^65,67–70^. Our data suggest a role for *JAG1*, a canonical Notch agonist, during cardiac mesoderm to cardiac progenitor cell fate specification. Interestingly, low efficiency CM differentiation showed increased IGF signaling at the −IWP2 stage, consistent with reports that IGF inhibition with LY294002 at this stage improves CM differentiation efficiency and introduction of insulin, an activator of IGF signaling inhibits CM differentiation^71–73^. At the +INS differentiation stage, we identified increases in PPAR signaling and glucose metabolism in high efficiency CM progenitors and increases in Wnt and MAPK signaling in low efficiency progenitors. Given proteomic and transcriptomic increases in glucose metabolism and PPAR signaling in high efficiency CM progenitors, we further investigated metabolomics and oxygen consumption and found that high efficiency CM progenitors displayed increased glycolytic and oxidative metabolism. Because we identified aberrant Wnt and MAPK signaling in low efficiency CM progenitors, we tested whether Wnt and MAPK inhibition could improve CM differentiation efficiency and found that Wnt or MAPK inhibition alone and in combination improved CM differentiation efficiency.

Molecular investigations to predict hPSC-CM differentiation efficiency during early differentiation stages are severely lacking^45,46,48,74^. Utilizing top DEGs from our multi-omic investigation, we designed an RT-qPCR panel aimed at predicting terminal CM purity at the +IWP2, −IWP2, and +INS stages. With this approach, we developed highly predictive machine learning models for terminal CM purity using just 4 genes at each timepoint and 11 genes in total. While some genes have known function during differentiation, several of these predictive genes have not been previously linked to CM differentiation. Notably, *LIN7A*, *RHOBTB3*, *ROBO3*, *SLC2A1*, and *S100A10* have relatively unknown roles in CM differentiation, although we have previously reported that *RHOBTB3* is predictive of CM differentiation efficiency at the cardiac progenitor stage^45^. *MESP1*^24^ and *SALL1*^75^ have been tied to mesoderm and lateral plate mesoderm development which are required stages prior to cardiomyocyte development. *ACKR3*, which was upregulated in low efficiency differentiations, has been linked to MAPK activation^76–78^. Furthermore, *DUSP5* was enriched in high efficiency CM differentiations and is known to inhibit MAPK signaling^79^. Surprisingly, *HRC* has been reported as a direct target of the MEF2 TF known to be involved in CM differentiation, yet our data detect *HRC* transcription in advance of *MEF2C* expression^80^. Interestingly, predictive gene panels outperformed canonical hPSC-CM differentiation genes at the +IWP2 and −IWP2 differentiation steps further highlighting the utility of investigations geared towards quantitatively predicting hPSC-CM differentiation efficiency during early differentiation stages.

Although bulk multi-omic analysis enabled deep investigation of divergent hPSC-CM differentiation outcomes, future studies employing single cell multi-omic approaches will be essential to resolve the precise lineage bifurcations and intermediate states that drive off-target cell fates^81^. Several reports have begun to define these transitions^51,82–87^, yet only a few have compared both high and low efficiency CM differentiations^21,46^. A recent study investigated the effect of CHIR concentration on mesoderm and CM differentiation and found that Wnt underactivation led to lack of differentiation or an epiblast-like gene signature while Wnt overactivation led to skeletal muscle differentiation^46^. Additionally, the mesendoderm transcription factor, *EOMES*, was highly correlated with successful cardiomyocyte differentiation^46^. Single cell multi-omic characterization approaches will facilitate the discovery of CM differentiation lineage branch points and enable signaling or transcription factor-based approaches to enable pure CM differentiation for therapeutic and modeling applications. Combining these approaches with our temporal bulk multi-omic roadmap could enable fully closed loop, feedback-controlled manufacturing systems for hPSC-CM cell products.

The multi-omic comparison of high and low efficiency CM differentiations reported here enabled comprehensive interrogation of the multi-omic landscape beyond the transcriptome. Notably, extracellular metabolomics identified non-invasive metabolites, such as glucose, suitable for real-time, in-line monitoring. Furthermore, intracellular metabolomics led us to further investigate metabolic cellular function, where oxygen consumption distinguished failed and successful differentiation batches during early differentiation stages, which could be easily measured to support next-generation manufacturing protocols. Integrating transcriptomic and proteomic findings allowed us to develop machine learning models of early differentiation success and implement protocol perturbations to enhance CM differentiation efficiency. With proper knowledge and manufacturing design, machine learning models based on -omic data could be utilized in tandem with targeted process interventions to improve differentiation success in a feedback-controlled manner. Together, these advances outline a multi-omic framework for reproducible manufacturing of complex cell products, such as hPSC-derived cells, accelerating high-quality cell therapies to treat a diverse spectrum of human diseases.

## Materials and Methods

### Human Pluripotent Stem Cell Maintenance

hPSCs (WTC11 hiPSCs, H9 hESCs, and IMR90-4 hiPSCs) were maintained as described previously^13,88^. hPSCs were cultured in mTeSR1 medium (STEMCELL Technologies 85850) on 6-well plates (Corning COSTAR 07-200-82) coated with Growth Factor Reduced Matrigel (0.5 mg/plate or 11 μg/cm^2^, Corning 354230). hPSCs were maintained in a cell culture incubator (Sanyo MCO-18AC; 37°C, 5% CO_2_, 95% RH). hPSCs were passaged with Versene (Life Technologies 15040066) every 3-5 days at 60-80% confluency.

### hPSC-Cardiomyocyte Differentiation

hPSCs were differentiated using the GiWi protocol as described previously^13,89^. On Day −2, hPSCs were detached and singularized with Accutase (Innovative Cell Technology AT104) for 5 min prior to quenching 1:1 v/v in DMEM/F12 (ThermoFisher 11330032). Subsequently, hPSCs were counted and resuspended at cell densities optimized for cardiomyocyte differentiation for each hPSC line. hPSCs were resuspended in mTeSR1 with 5 μM Y-27632 (Tocris 1254) and seeded onto Matrigel-coated 12-well plates in 2 mL of media. To facilitate even cell attachment on Day −2, plates were left at room temperature for 30 min prior to maintenance in a cell culture incubator. On Day −1, medium was replaced with mTeSR1. On Day 0, differentiation was induced with CHIR99021 (Selleckchem S1263) at a concentration between 6-12 μM, optimized for cardiomyocyte differentiation for each hPSC line. CHIR99021 was added to RPMI1640 medium (Life Technologies 11875119) with 2% v/v B27 minus insulin (B27-). After Day 0, protocol timing is dependent on hPSC line as previously established^36,45,88^. For WTC11 hiPSCs, media was replaced with B27- and 5 μM IWP2 (Tocris 3533) on Day 2 and on Day 4 with B27-. On Days 6, 8, 10, and 13, media was replaced with RPMI1640 with 2% v/v B27 with insulin (B27+, Life Technologies 17504044) until collection on Day 16. In GiWi hPSCs (IMR90-4 hiPSCs and H9 hESCs), media was replaced on Days 1 and 5 with B27- and on Day 3 with 1:1 v/v B27-:conditioned medium with 5 μM IWP2. On Days 7, 10, and 13, media was replaced with B27+ until collection on Day 16.

### hPSC-Cardiomyocyte Differentiation Rescue with Wnt and MAPK Inhibition

WTC11 hiPSCs were differentiated to D5 cardiac progenitor cells (CPCs) as described above. CPCs were cryopreserved prior to thawing and replating as previously described^36^. Cells were lifted with Accutase and resuspended in 6:3:1 v/v/v B27-, fetal bovine serum (FBS, R&D Systems S12450), and dimethyl sulfoxide (DMSO, Sigma-Aldrich D2650) prior to controlled-rate freezing at −1°C per min in a Mr. Frosty Freezing Container (ThermoFisher 51000001). Thawed CPCs were reseeded for hPSC-CM differentiation at a surface area ratio between 1:1-1:2.5. CPCs were treated with small molecules IWP2 (5 µM) and/or FHPI (VWR 103538-448, 10 µM) in B27+ medium from Day 5-8. Small molecules were added on thaw on Day 5 and on Day 6 during the B27+media change. Control samples were treated with DMSO as a vehicle control. Subsequently, cardiomyocyte differentiation was carried out as described above prior to Day 16 collection for flow cytometry to assess CM purity and number.

### Multi-omic Sample Collection Design

On media exchange days throughout the differentiation, wells from each WTC11 hiPSC-CM differentiation batch were sacrificed for collection and subsequent analysis. This included Days 0, 2, 4, 6, 8, 10, 13, and 16. For RNA collection, a set of wells was rinsed once with DPBS prior to 1 min incubation with 500 µL cold Trizol reagent (ThermoFisher 15596018). Subsequently, the Trizol-cell mixture was scraped into 1.5 mL microcentrifuge tubes, snap-frozen in liquid nitrogen, and stored at −80°C (RNA). For simultaneous protein, intracellular metabolite, and extracellular metabolite collection from a parallel set of wells (parallel to RNA wells), 1 mL of media was collected and transferred to a fresh microcentrifuge tube, snap-frozen in liquid nitrogen, and stored at −80°C (extracellular metabolites). Cells from the same well as media collection for extracellular metabolomics were collected with cold (4°C) 100% methanol for sequential extraction of intracellular metabolites and proteins for mass spectrometry per our previously published methods as described below.^90^ Each sample consisted of 1 well of a 12-well plate, which contained approximately 1-2 million cells. All -omic collections were completed prior to media change and small molecule addition (if applicable) on the day of collection.

### Nuclear Magnetic Resonance (NMR) Extracellular Metabolomics

Conditioned cell culture media was snap-frozen in liquid nitrogen and stored at −80°C until analysis. Samples were subsequently prepared in a method adapted from Grouw et al.^91^ Media samples were thawed at 4°C and centrifuged at 20 g for 15 min to pellet insoluble debris. Supernatant was transferred to fresh microcentrifuge tubes and supplemented with 10% v/v NMR prep solution (3.333 mM DSS-d6 in D_2_O). 550 uL was loaded into a fresh 5 mm NMR tube. Remaining supernatants from each sample were pooled as an internal pooled control standard. Controls lacking cells were also collected, which were subjected to the same conditions in the absence of cells. Samples were run on the following pulse programs on a Bruker Avance III HD, 600 MHz NMR equipped with a 5 mm cryoprobe QXI (1H/19F/13C/15N) Z-axis gradient with the following pulse programs: NOESYPR1D, JRES, and HSQC. 1D ^1^H NOESYPR1D files were processed with the Edison Lab’s Metabolomics Workbench program in MATLAB for spectral processing, peak binning, and AUC quantitation.^92^ These AUC values were analyzed using the statistical analysis module of MetaboAnalyst for statistical comparisons of interest and used in multi-omic factor analysis described below^43^. Subsequently, metabolite identifications were confirmed from 2D HSQC files for 25 extracellular metabolites from Day 0-6 and concentrations were quantified using Chenomx NMR Suite v10.1. Extracellular metabolite concentration data were normalized by median and log-transformed prior to statistical analysis using MetaboAnalyst v6.0^43,93^. At each timepoint, Benjamini-Hochberg FDR adjusted p-values from unpaired t-tests were used to determine significant differences between groups. Principal component analyses, volcano plots, and quantitative enrichment analyses were used to visualize differences between groups.

### Sequential Extraction of Intracellular Metabolites and Proteins

After cell culture media was collected for NMR-based extracellular metabolomics, intracellular metabolites and proteins were sequentially extracted from the same well using a single step methanol-based quench as previously described^90,94^. Prior to quenching, wells were rinsed twice with DPBS (calcium/magnesium free). To quench metabolic activity, cells were treated for 1 min with 1 mL of cold (4°C) 100% methanol. Subsequently, the methanol-cell mixture was scraped into microcentrifuge tubes, vortexed for 10 s, and mixed at 4°C for 10 min. To create a metabolite supernatant and protein cell pellet, the methanol-cell mixture was centrifuged for 5 min at 14,000*g*. The protein cell pellet was washed with 200 µL of DPBS. Intracellular metabolite and protein samples were vacuum-dried and stored at −80°C.

### Untargeted Mass Spectrometry-based Intracellular Metabolomics

For sample preparation, intracellular metabolite samples were thawed on ice prior to rehydration with cold 50% methanol and mass spectrometry analysis by flow injection to a Bruker 12T solariX Fourier transform ion cyclotron resonance (FTICR) mass spectrometer as previously described^90^. 50% methanol metabolite reconstitution was performed stepwise by addition of 100 µL of 100% methanol with vortexing followed by addition of 100 µL mass spectrometry-grade water with vortexing.

For data acquisition, flow-injection electrospray Fourier transform ion cyclotron resonance (FIE-FTICR) mass spectrometry-based metabolomics was conducted on a Waters ACQUITY UPLC M-Class System (Waters Corporation) coupled to a Bruker solariX 12T FTICR mass spectrometer (Bruker Daltonics) operated with no LC column using previously established acquisition parameters^95,96^. Samples were introduced into the mass spectrometer via electrospray ionization. Mass spectrometry-grade solvents were used for mobile phase preparation for positive (0.1% FA in 50% methanol in water) and negative (10 mM ammonium acetate in 50% methanol in water) ionization modes. A 5-min isocratic flow of 100% pre-mixed mobile phase was delivered at 15 μL/min, with 5 μL injection volume per sample. In brief, 50 scans per spectrum were acquired with an ion accumulation time of 0.1 s per spectrum, resulting in a total data collection time of 3.5 min per sample with a 1.5 min wash step. Prior to data acquisition, the FTICR mass spectrometer was calibrated in each ionization mode using 1 mM NaTFA.

For metabolite feature detection, intracellular metabolomics data were processed in MetaboScape 2021b (Bruker Daltonics). Bucket lists for data acquired in positive and negative ionization modes were generated using the T-Rex 2D algorithm and subsequently merged into a single combined list using a mass error tolerance <1.0 ppm. To remove background signals, a blank filtering step was applied, excluding features with intensities less than three times that observed in the blank sample. SmartFormula annotations were carried out in MetaboScape^96^. Metabolic features were annotated based on accurate mass matching against the Mass Bank of North America and the Human Metabolome Database^97^, using a mass error tolerance ≤3 ppm. KEGG, HMDB IDs, and PubChem IDs were assigned to each identified metabolite. Intensities were filtered using an intensity minimum of 8E6 prior to statistical analysis. All annotated metabolite features were manually confirmed in DataAnalysis software (Bruker Daltonics) to validate feature detection in MetaboScape. Metabolite data were treated using MetaboAnalyst by estimating missing values by the median value of feature intensities in each sample, normalized by sum, and log-transformed prior to statistical analysis. At each timepoint, Benjamini-Hochberg FDR adjusted p-values from unpaired t-tests were used to determine significant differences between groups. Principal component analyses and quantitative enrichment analyses were used to visualize differences between groups.

### Global Mass Spectrometry-based Bottom-Up Proteomics

For sample preparation, the protein-rich dried cell pellet was solubilized in 10 μL of buffer containing 0.25% w/v Azo^95,98^. Samples were subjected to sonication and multiple freeze-thaw cycles to facilitate protein solubilization, then diluted to 0.05% w/v Azo. Protein concentrations were normalized using the Bradford assay. Samples were reduced with 10 mM DTT for 1 h at 37°C and alkylated with 20 mM IAA for 30 min at room temperature in dark, followed by enzymatic digestion using Trypsin Gold (Promega) at a 50:1 protein-to-enzyme ratio for over 3 h at 37°C. Enzymatic activity was quenched with formic acid, and samples were subsequently irradiated at 305 nm for 5 min to degrade Azo. Peptides were desalted with C18 ZipTips, dried under vacuum, and rehydrated in 0.1% formic acid in water at a concentration of 0.2 mg/mL. Peptide concentrations were quantified using a NanoDrop 2000 prior to LC–MS analysis.

For data acquisition, Tryptic peptide analyses were carried out using a timsTOF Pro mass spectrometer (Bruker Daltonics) connected to a nanoElute UHPLC system (Bruker Daltonics, Bremen) equipped with a Captivespray nano-electrospray ion source. For each run, approximately 200 ng of peptide material was injected onto an IonOpticks C18 capillary column (25 cm×75 μm i.d., 1.7μm particles). Peptide separation occurred at 55°C with a flow rate of 400 nL/min under a stepwise gradient: 2–17% solvent B over 60 min, followed by 17–25% B from 60–90 min, and a final 30 min column wash at 85% B. Solvent A was 0.1% formic acid in water, and solvent B was 0.1% formic acid in acetonitrile. The mass spectrometer operated in Parallel Accumulation–Serial Fragmentation (PASEF) mode^99^, acquiring MS and MS/MS data across an m/z window of 100–1700. Ion mobility separation ranged from 0.60 to 1.60 V·s/cm² with a 100 ms ramp time. Data-dependent acquisition (DDA) included 10 PASEF MS/MS events per cycle using a locked duty cycle of 100%. Precursors exceeding 20,000 intensity units were excluded for 0.4 min and automatically reconsidered after 4 min. Collision-induced dissociation (CID) energies were adjusted according to ion mobility (1/K₀). Peptide loading and injection consistency were confirmed by comparing total ion current intensities to a 200 ng human K562 whole-cell lysate standard (Promega) prepared at 0.2 µg/µL.

Protein identification was performed using MaxQuant v1.6.17.0 against the UniProt human database with a 1% false discovery rate for proteins and peptides^100^. TIMS-DDA runs used a half-width of 6, resolution of 32,000, and precursor mass tolerance ≤30 ppm. Peptides required ≥7 amino acids with a fixed modification for carbamidomethylation and variable modifications for methionine oxidation and N-terminal acetylation. Enzyme specificity was set to trypsin/P with 2 missed cleavages allowed. Label free quantification (LFQ) used a minimum ratio count of 2 with classic normalization. Match between runs employed a maximum retention window of 0.7 min and the ion mobility window was set to 0.05 V·s/cm. MS/MS spectra were required for LFQ comparisons.

Proteomics data were processed using ProStar and DAPAR R packages^101^. LFQ intensities were log2-transformed. Proteins absent from at least 2 of 3 or 3 of 4 biological replicates in all groups were removed, along with contaminants and reverse hits. Missing values were imputed using the slsa algorithm for partial data and 2.5% of the observed intensity quantile for proteins absent in one condition. Subsequent analyses were performed with MetaboAnalyst and R packages as described for metabolomics and transcriptomics.

### RNA Extraction, Sequencing, and Analysis

RNA was extracted from Trizol-collected samples via tri-phasic separation, then purified and concentrated using Zymo Direct-zol MiniPrep Plus columns (Zymo R2072) with on-column DNAse treatment per kit instructions. Purified RNA concentration was measured with a NanoDrop spectrophotometer (ThermoFisher ND2000c) and stored at −80°C. RNA sample quality and quantity were assessed for 18S and 28S ribosomal RNA presence using A260/A280 (∼2.0) and A260/A230 (>2.0) ratios. Prior to library construction and sequencing, RNA was quantified using Qubit on an Agilent 2100 Bioanalyzer. Libraries were built from 500 ng total RNA per sample using the Takara SMARTer Stranded Total RNA Kit - HI Mammalian and sequenced on an Illumina NovaSeq 6000.

Raw FASTQ files were aligned to the human genome (hg38+decoy) using STAR v2.5.3a^102^. Gene-level counts were generated with featureCounts v2.0.3^103^. Differential expression analysis was performed in R v4.1.3^104^ using DESeq2, with PCA plots generated from variance-stabilized counts. Differentially expressed genes were defined by log₂ fold change>1 and padj<0.05. QuaternaryProd analysis was conducted using v1.28.0^40^. Overrepresentation analyses, gene set enrichment analyses, upset plots and corresponding histograms were created with clusterProfiler v4.0^39^. Multi-omic factor analysis was performed using MOFA2 v0.6.7 using the MEFISTO framework and the top 5000 most variable transcripts^105^. Fgsea v1.26.0 was used for gene set co-regulation analysis (GESECA)^38^. For multi-omic gene set enrichment analysis, multiGSEA v1.16.0 was used^106^. PACNet analysis was performed using the online tool^37^. Custom scripts were used to create heatmaps, volcano plots, and parity plots. Protein-protein interaction diagrams were created with STRING^107^.

### Public RNA Sequencing Dataset Analysis

The timecourse RNAseq FASTQ files deposited by Strober et al. (GSE122380)^20^ were acquired from the Gene Expression Omnibus (GEO)^108^ and processed the same as the RNAseq data described above.

### Flow Cytometry

Cardiomyocyte purity was assessed on D16 via flow cytometry for cardiac troponin T (cTnT) as described in prior work^13^. Accutase was used for cell singularization, 1% paraformaldehyde was used for fixation, and 90% methanol was used for −20°C storage until analysis. Cells were rinsed in flow buffer (FB = 0.5% w/v BSA in Dulbecco′s Phosphate Buffered Saline, DPBS) and treated overnight at 4°C in 1:1000 msIgG1-anti-cTnT primary antibody (ThermoFisher MA5-12960) in FB + 0.1% v/v Triton X-100. After rinsing with FB the subsequent day, the cells were treated in the dark at room temperature for 30-60 min in either 1:1000 AF488-anti-msIgG1 secondary antibody (ThermoFisher A-21121) or 1:1000 AF647-anti-msIgG1 secondary antibody (ThermoFisher A21240) in FB + 0.1% v/v Triton X-100. Cells were rinsed and resuspended in 200-500 µL FB for analysis on a BD Accuri C6 Plus or ThermoFisher Attune flow cytometer. Undifferentiated hPSC negative control samples were stained in parallel, and gating was performed using BD Accuri C6 Plus or FlowJo software.

### Immunocytochemistry

Day 16 adherent cells were rinsed with DPBS and fixed with 1% paraformaldehyde (PFA) in DPBS for 30 min at room temperature. Wells were rinsed twice with DPBS, blocked with 1% BSA in DPBS, and stored at 4°C until immunostaining, which proceeded as described for flow cytometry. Following secondary antibody incubation, wells were stained with 2 μg/mL Hoechst 33342 (Invitrogen H3570) in DPBS for 5 min at room temperature in the dark, rinsed twice with DBS, and immersed in DBPS for imaging. Images were acquired at indicated magnifications on a Nikon Ti2e with an Aura III light engine (Lumencor 80-10306) and ORCA-Flash4.0 camera (Hamamatsu C13440-20CU).

### Seahorse MitoStress Test

Mitochondrial respiration in hPSC-CPCs (+INS differentiation step, D6 in WTC11 hiPSCs) was evaluated using a Seahorse XF96 extracellular flux analyzer. Cells were lifted on D5, cryopreserved, and replated onto Matrigel-coated XF96 cell culture microplates at a surface area ratio of 1:2.5 as previously described^36^. Following a one-day recovery, the MitoStress Test assay was performed. On D6, cells were equilibrated for 1 hr at 37°C (non-CO₂) in Seahorse RPMI basal medium with 5 mM glucose, 1 mM sodium pyruvate, and 2 mM glutamine. Mitochondrial stress was stimulated via subsequent injections of 1.5 μM oligomycin (ATP synthase inhibitor), 1 μM FCCP (uncoupler), and 0.5 μM rotenone/antimycin A (Complex I/III inhibitors). Cell numbers were normalized using Hoechst 33342 (10x magnification). One high and low efficiency differentiation batch of cardiomyocyte purity-validated cardiac progenitors were assayed for oxygen consumption rate (OCR) with n=46 and n=30 well replicates respectively. OCR calculations proceeded as follows: basal OCR (average baseline, timepoints 1–3); maximal OCR (peak FCCP-stimulated minus residual post-rotenone/antimycin A); ATP-linked OCR (basal minus residual post oligomycin); Non-mitochondrial OCR (residual post-rotenone/antimycin A).

### Mitochondrial/Nuclear DNA Content Analysis

Genomic DNA was isolated from Trizol samples via tri-phasic extraction. Mitochondrial (mito) and nuclear (nuc) genomic DNA content were quantified by qPCR as previously described^109^. Mitochondrial DNA content was normalized to nuclear DNA, and resulting ratios were normalized to values from low efficiency hPSC-CM progenitors at each differentiation step. Primer sequences are listed in Table S1.

### Reverse Transcription, Quantitative Polymerase Chain Reaction (RT-qPCR), Primer Design

Total RNA (2 µg) was reverse transcribed using the Qiagen Omniscript RT Kit (205113) with RNaseOUT (Life Technologies 10777-019) and Oligo dT(20) primers (Life Technologies 18418020) per manufacturer instructions. qPCR was performed with 10ng cDNA per reaction in a 25 µL volume containing 12.5 µL PowerUp SYBR Green Master Mix (ThermoFisher A25780), 0.125 µL of 100 µM primers (IDT), and nuclease-free water. Reactions were run on an AriaMx Real-Time PCR System (Agilent G8830A) for 40 cycles (95°C, 15 s; 60°C, 60 s), followed by melt-curve analysis to confirm single-product amplification. The geometric mean of *ZNF384*, *DDB1*, and *EDF1* was used as the reference gene as described previously^88^. Primers were designed using NCBI Primer-BLAST, and sequences were ordered from IDT (Table S1). Amplicon size was confirmed by agarose gel electrophoresis. Primers producing a single, correctly sized band in positive control cDNA were assessed for qPCR efficiency using a four-point 1:5 dilution series ([cDNA] = 10, 2, 0.4, 0.08 ng/reaction) and a no template control ([cDNA] = 0 ng/reaction). Primers showing no NTC amplification and efficiencies of 90–110% were approved for further use.

### Generation of RT-qPCR Models

RT-qPCR analysis of 54 genes with potential cardiomyocyte-predictive power, including canonical markers, was performed across 22 additional differentiation batches from H9 hESCs, IMR90-4 hiPSCs, and WTC11 hiPSCs. Gene expression was measured at +CHIR, +IWP2, –IWP2, and +INS stages. Partial least squares regression (PLSR) models were generated using MetaboAnalystR v3.30 and the pls v2.8-3 package. Step-specific PLSR models used 5 components with leave-one-out cross-validation. The top 25 genes correlated with terminal CM purity at each stage were plotted by correlation coefficient, controlling for cell line. For nonlinear model evolution, DataModeler v9.7 (Evolved Analytics) was used to generate step-specific models and models from 10 canonical CM differentiation markers (Tables S2-S9). Step-specific models were evolved over multiple rounds to reduce gene sets, while canonical models were evolved for one round to obtain comparable gene numbers. Model performance was compared by root mean squared error.

### Statistical Analysis

Multi-omics data (Figure 1) were collected from 3–4 wells within one WTC11 hiPSC-CM differentiation per condition (high vs. low efficiency). Confirmatory RNA-seq (Figure 2) used one well per timepoint across six differentiations including high and low efficiency hPSC-CM differentiations for three lines (H9 hESCs, IMR90-4 hiPSCs, WTC11 hiPSCs). Statistical tests are provided in figure legends. Annotations include: ∗ (p<0.05), ∗∗ (p<0.01), ∗∗∗ (p<0.001), ∗∗∗∗ (p<0.0001), and ns (p>0.05).

## Acknowledgments

The authors thank the University of Wisconsin Carbone Cancer Center Flow Cytometry Laboratory, supported by P30 CA014520, for use of its facilities and services. The authors also would like to acknowledge the UW-Madison Human Proteomics Program Mass Spectrometry Facility (initially funded by the Wisconsin partnership funds) in obtaining mass spectrometry data and NIH S10OD018475 for the acquisition of ultra-high resolution mass spectrometer for biomedical research. This study made use of the National Magnetic Resonance Facility at Madison, which is supported by NIH grant R24GM141526. Helium recovery equipment, computers, and infrastructure for data archive were funded by the University of Wisconsin–Madison, NIH R24GM141526, and National Science Foundation NSF 1946970 (NSF Mid-Scale Research Infrastructure Big Idea). This work was completed in part with resources provided by the University of Wisconsin Biotechnology Center- Gene Expression Center (RRID: SCR_017757).

## Author contributions

Conceptualization: AKF, ADS, SPP

Data Curation: AKF, ADS, EFB, MRP

Formal Analysis: AKF, ADS, EFB, MRP

Funding Acquisition: TJK, YG, SPP

Investigation: AKF, ADS, EFB, YZ, TJK, YG, SPP

Methodology: AKF, ADS, EFB, YG, SPP

Project Administration: TJK, YG, SPP

Resources: YG, SPP

Software: AKF, ADS

Supervision: TJK, YG, SPP

Validation: AKF, ADS, EFB, YG, SPP

Writing—original draft: AKF, ADS

Writing—review & editing: All authors

## Data and materials availability

All data needed to evaluate the conclusions in the paper are present in the paper and/or the Supplementary Materials.

Next-generation RNA sequencing data has been deposited on the NCBI Gene Expression Omnibus^108^ under accession number GSE314132.

The mass spectrometry proteomics data have been deposited to the ProteomeXchange Consortium via the PRIDE^110^ partner repository with the dataset identifier PXD072237.

The intracellular mass spectrometry metabolomics data has been deposited to Zenodo and is available under the following DOI: doi.org/10.5281/zenodo.17992183.

Any additional requests for data can submitted to Sean Palecek.

