## Supplemental Figures and Tables for "Early multi-omic signatures and machine learning models predict cardiomyocyte differentiation efficiency and enable robust hPSC differentiation to cardiomyocytes"

**This PDF file includes:**

Figs. S1 to S8
Tables S1 to S9

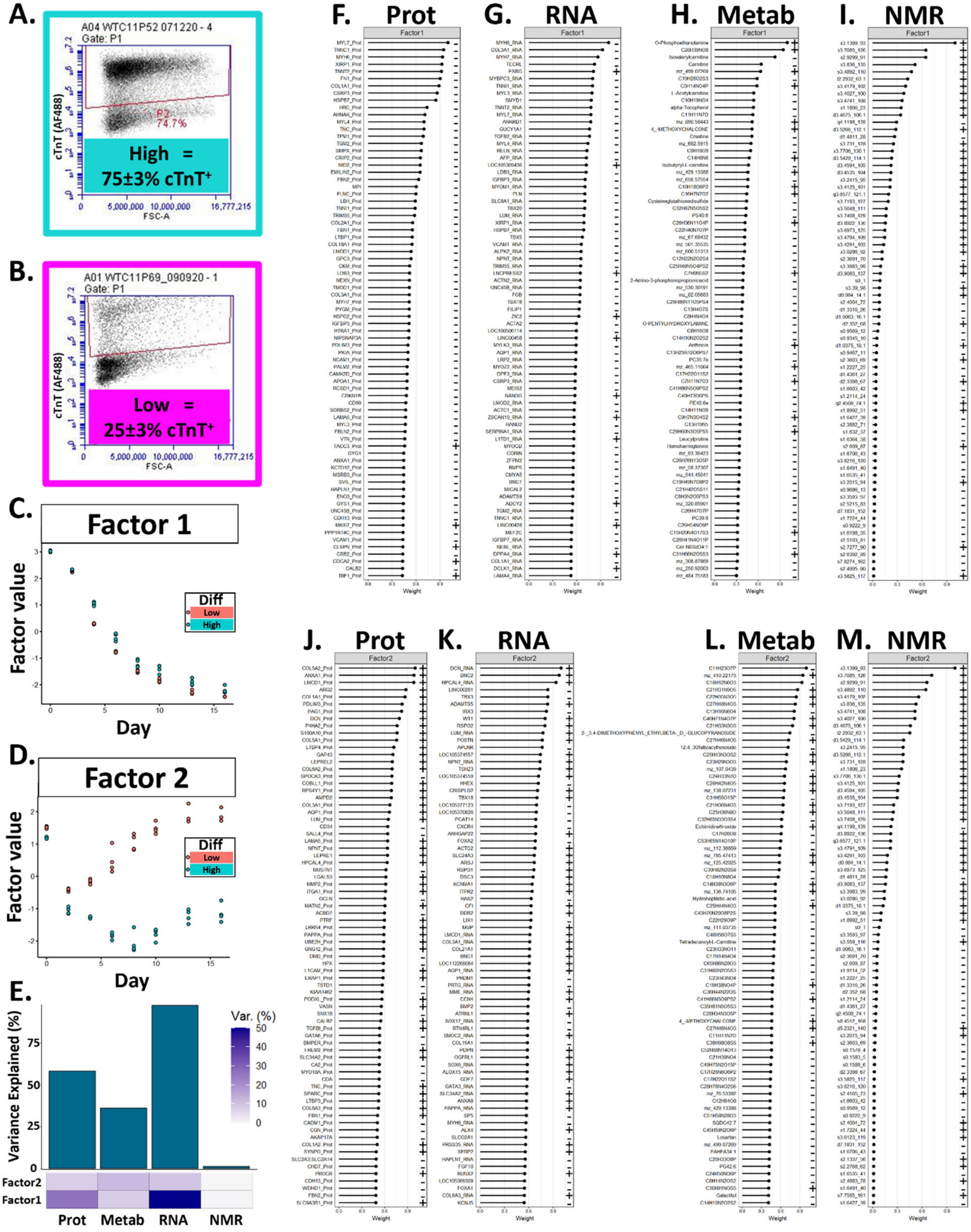


**Figure S1. Representative flow cytometry gating and Multi-omic Factor Analysis (MOFA) factor values and feature weights. A-B)** Representative flow cytometry dot plots for Day 16 hPSC-CMs for **A)** high (turquoise) and **B)** low (magenta) differentiation efficiency batches. **C-**

**D)** Overall MOFA factor values for high (turquoise) and low (red) differentiation efficiency batches for **C)** Factor 1 and **D)** Factor 2 for each group at each timepoint. **E)** Percent variance explained by each -ome for each factor. **F-I)** Factor 1 feature weights for the top 50 features for the **F)** proteome, **G)** transcriptome, **H)** intracellular metabolome, and **I)** extracellular metabolome. Each feature has a positive or negative feature weight (symbol on right side of plot) indicating the contribution to the calculation of overall factor value. **J-M)** Factor 2 feature weights for the top 50 features for the **J)** proteome, **K)** transcriptome, **L)** intracellular metabolome, and **M)** extracellular metabolome. Each feature has a positive or negative feature weight (symbol on right side of plot) indicating the contribution to the calculation of overall factor value.

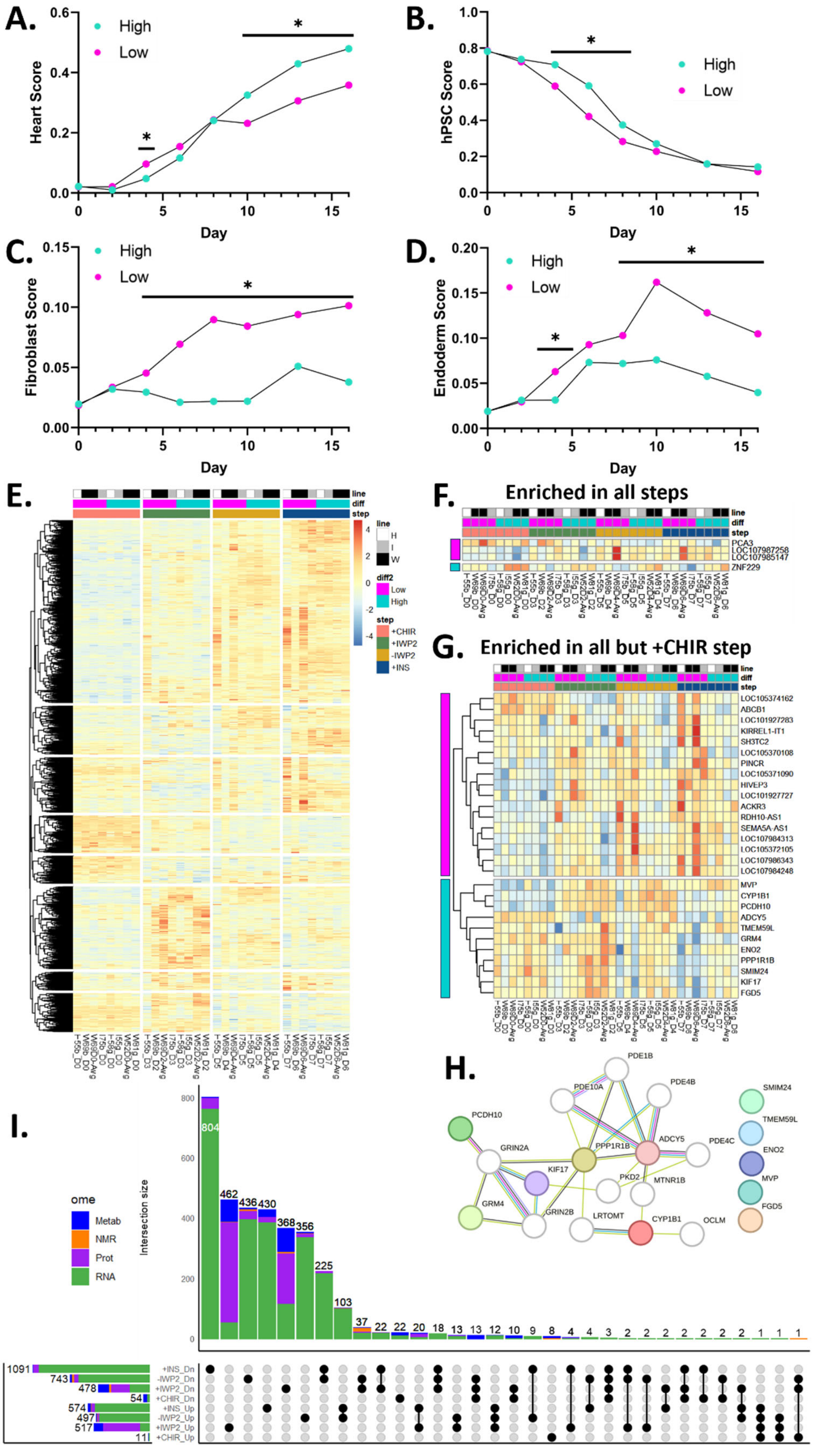
A.
Heart Score
High
Low
Day
B.
hPSC Score
C.
Fibroblast Score
D.
Endoderm Score
E.
line
diff
step
diff2
Low
High
+CHIR
+IWP2
-IWP2
+INS
F.
Enriched in all steps
PCA3
LOC107987258
LOC107985147
ZNF229
G.
Enriched in all but +CHIR step
LOC105374162
ABCB1
LOC101927283
KIRREL1-IT1
SH3TC2
LOC105370108
PINCR
LOC105371090
HIVEP3
LOC101927727
ACKR3
RDH10-AS1
SEMA5A-AS1
LOC107984313
LOC105372105
LOC107986343
LOC107984248
MVP
CYP1B1
PCDH10
ADCY5
TMEM59L
GRM4
ENO2
PPP1R1B
SMIM24
KIF17
FGD5
H.
PDE1B
PDE10A
PDE4B
PCDH10
GRIN2A
PPP1R1B
ADCY5
PDE4C
KIF17
PKD2
MTNR1B
GRM4
GRIN2B
LRTOMT
CYP1B1
OCLM
SMIM24
TMEM59L
ENO2
MVP
FGD5
I.
ome
Metab
NMR
Prot
RNA
Intersection size
+INS_Dn
-IWP2_Dn
+IWP2_Dn
+CHIR_Dn
+INS_Up
-IWP2_Up
+IWP2_Up
+CHIR_Up

**Figure S2. PACNet classification scores and differential transcripts are conserved across hPSC lines and differentiation steps. A-D)** PACNet classification scores (range: 0-1) comparing high and low efficiency groups across the differentiation time course for several cell or tissue types. **A)** Heart classification scores. **B)** Human pluripotent stem cell (hPSC) classification scores. **C)** Fibroblast classification scores. **D)** Endoderm classification scores calculated from the addition of lung, liver, and intestine classification scores. P-values from unpaired t-tests. **E)** Heatmap of the highest variance transcripts across all tested hPSC lines (H9, IMR90-4, and WTC11) and differentiation steps (TPM). **F)** Heatmap of the four transcripts consistently differentially expressed across all differentiation steps (TPM). Transcripts enriched in low efficiency differentiation batches are labeled with the vertical magenta box. The single transcript enriched in the high efficiency differentiation batches is labeled with the vertical turquoise box. **G)** Heatmap of the 28 transcripts consistently differentially expressed across all differentiation steps except the +CHIR step (TPM). Transcripts enriched in low efficiency differentiation batches are labeled with the vertical magenta box. Transcripts enriched in the high efficiency differentiation batches are labeled with the vertical turquoise box. **H)** STRING network of the 11 genes conserved as enriched in the high efficiency differentiation batches (bottom of panel d coinciding with the vertical turquoise box). **I)** UpSet plot of differential features at each differentiation step (as in Figure 2H) further separated by direction of change where Up corresponds to enriched in high efficiency differentiation batches and Dn (Down) corresponds to enriched in low efficiency differentiation batches.

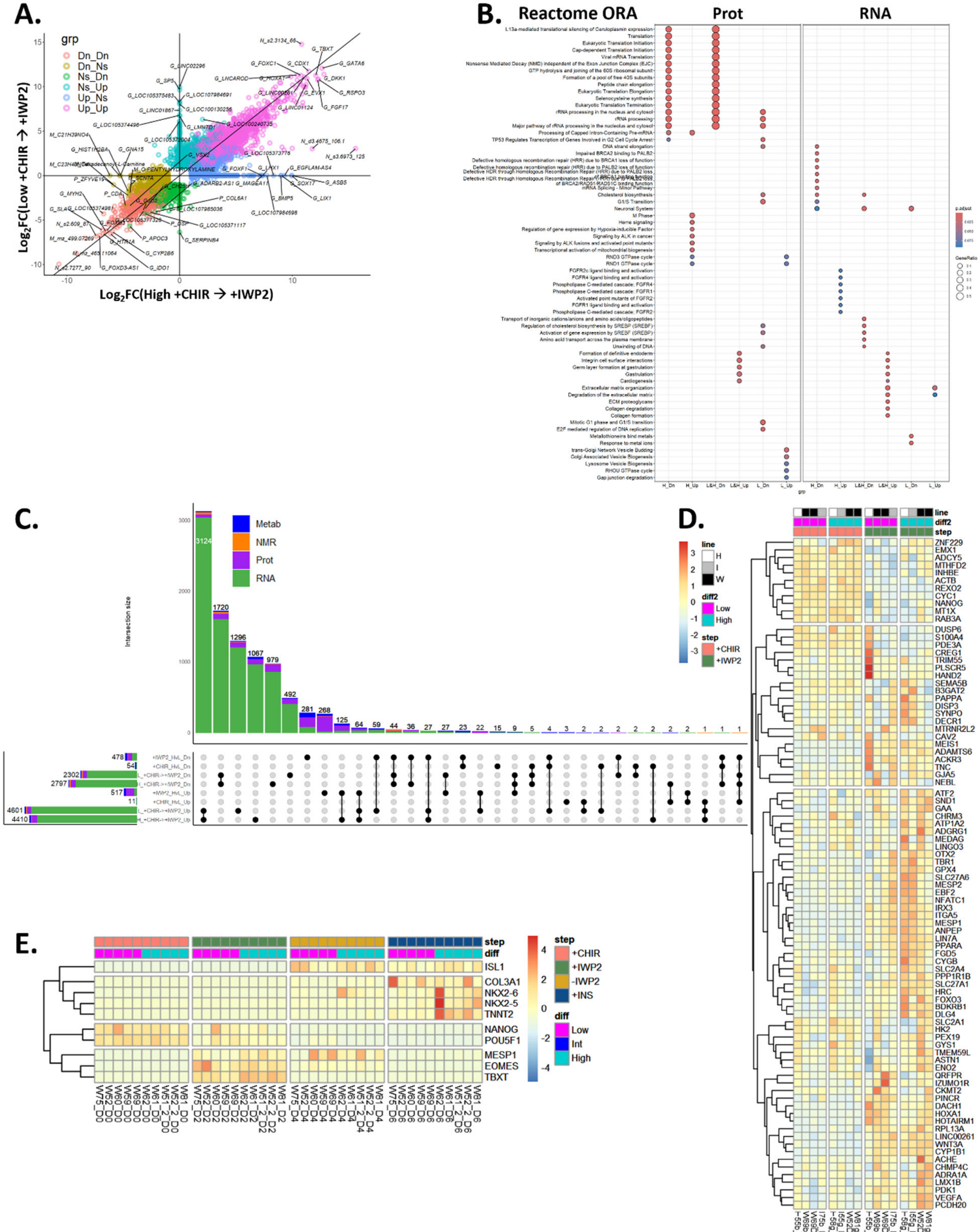

**Figure S3. Comparison of proteomic and transcriptomic pathway analysis and differential feature abundance for the +CHIR to +IWP2 transition. A)** Parity plot comparing the differential features changing from the +CHIR step to the +IWP2 step in high (x-axis) and low (y-axis) efficiency differentiations. Prefixes for each -omic feature are G = gene, P = protein, M = intracellular metabolite, N = extracellular metabolite. **B)** Overrepresentation analysis for differentially expressed proteomic and transcriptomic features for the Reactome database. Comparisons are denoted as H (high efficiency), L (low efficiency), Up (enriched), and Dn (decreased) on the x-axis. Comparisons shown are enriched during the +CHIR to +IWP2 transition in either high, low, or both differentiation efficiency groups as indicated. **C)** UpSet plot of differential features for each comparison (as in Figure 3G) further separated by direction of change where Up corresponds to enriched and Dn (Down) corresponds to decreased for temporal or high (H) versus low (L) differentiation efficiency comparisons as indicated. **E)** Heatmap of transcriptomic data for 88 of the top differentially expressed transcripts at the +CHIR and +IWP2 differentiation step conserved across 3 hPSC lines (TPM). **F)** Heatmap of RT-qPCR data for 10 canonical stage-specific markers throughout the hPSC-CM differentiation for 10 additional differentiation batches from WTC11 hiPSCs across all 4 differentiation steps (2^-ΔCT). ΔCT = cycle threshold of gene minus cycle threshold of the geomean of 3 reference genes (*ZNF384*, *EDF1*, *DDB1*). Line = cell line (H = H9, I = IMR90-4, W = WTC11).

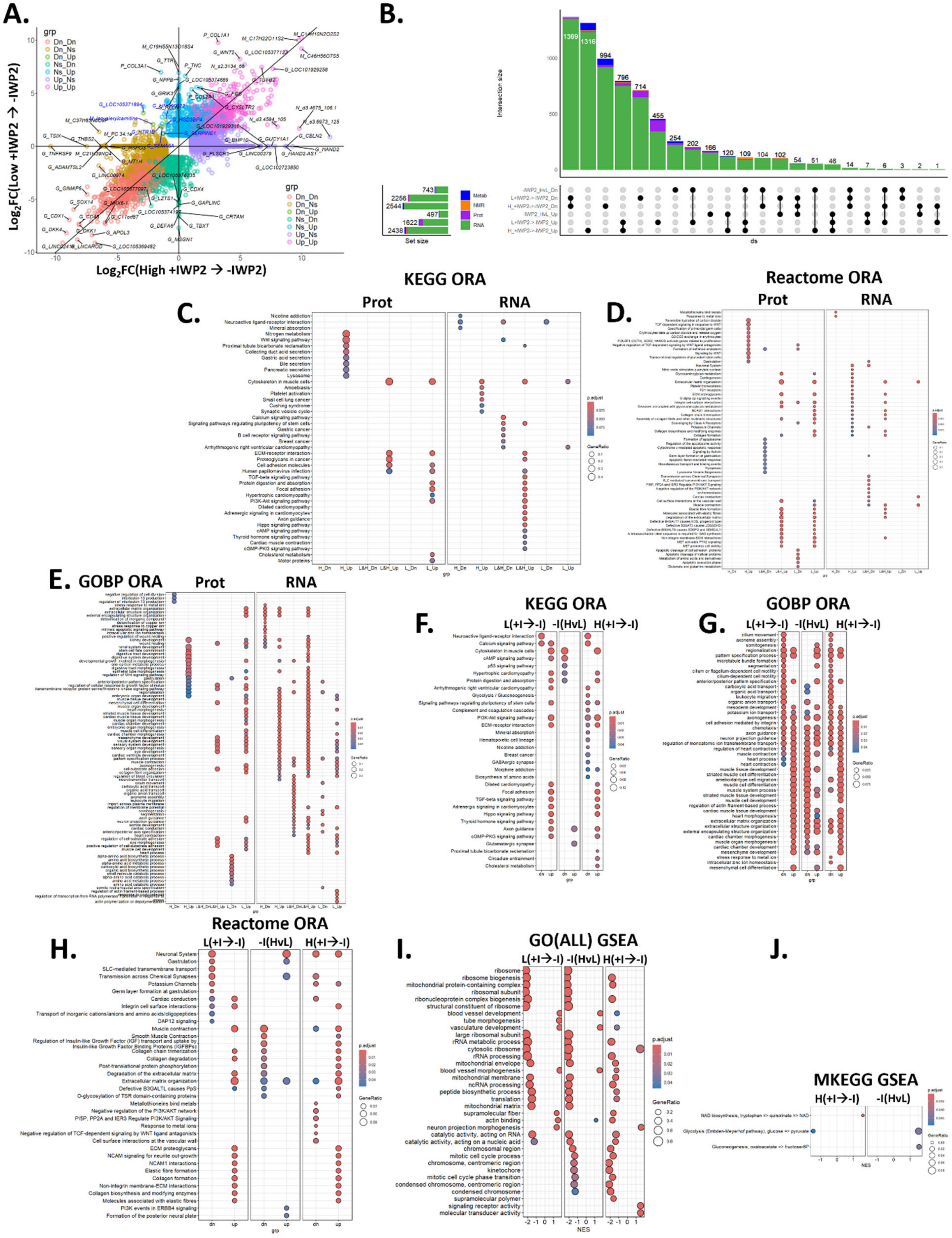


**Figure S4. Differential feature abundance, comparison of proteomic and transcriptomic pathway analysis, and additional transcriptomic pathway analysis for the +IWP2 to -IWP2 transition. A)** Parity plot comparing the differential features changing from the +IWP2 step to the -IWP2 step in high (x-axis) and low (y-axis) efficiency differentiations. **B)** UpSet plot of differential features for each comparison (as in Figure 4B) further separated by direction of change where Up corresponds to enriched and Dn (Down) corresponds to decreased for the temporal or high (H) versus low (L) differentiation efficiencies as indicated. **C-E)** Overrepresentation analysis for differentially expressed proteomic and transcriptomic features for the **C)** KEGG, **D)** Reactome, **E)** GO Biological Process databases. Comparisons are denoted as H (high efficiency), L (low efficiency), Up (enriched), and Dn (decreased) on the x-axis. Comparisons shown are enriched during the +IWP2 to -IWP2 transition in either high, low, or both differentiation efficiency groups as indicated. **F-H)** Overrepresentation analysis for differentially expressed transcriptomic features for the **F)** KEGG, **G)** GO Biological Process, **H)** Reactome databases. Comparisons are shown for temporal or high (H) versus low (L) differentiation efficiencies as indicated. **I-J)** Gene set enrichment analysis for differentially expressed transcriptomic features for the **I)** GO and **J)** Metabolic KEGG databases. Comparisons are shown for temporal or high (H) versus low (L) differentiation efficiencies as indicated.

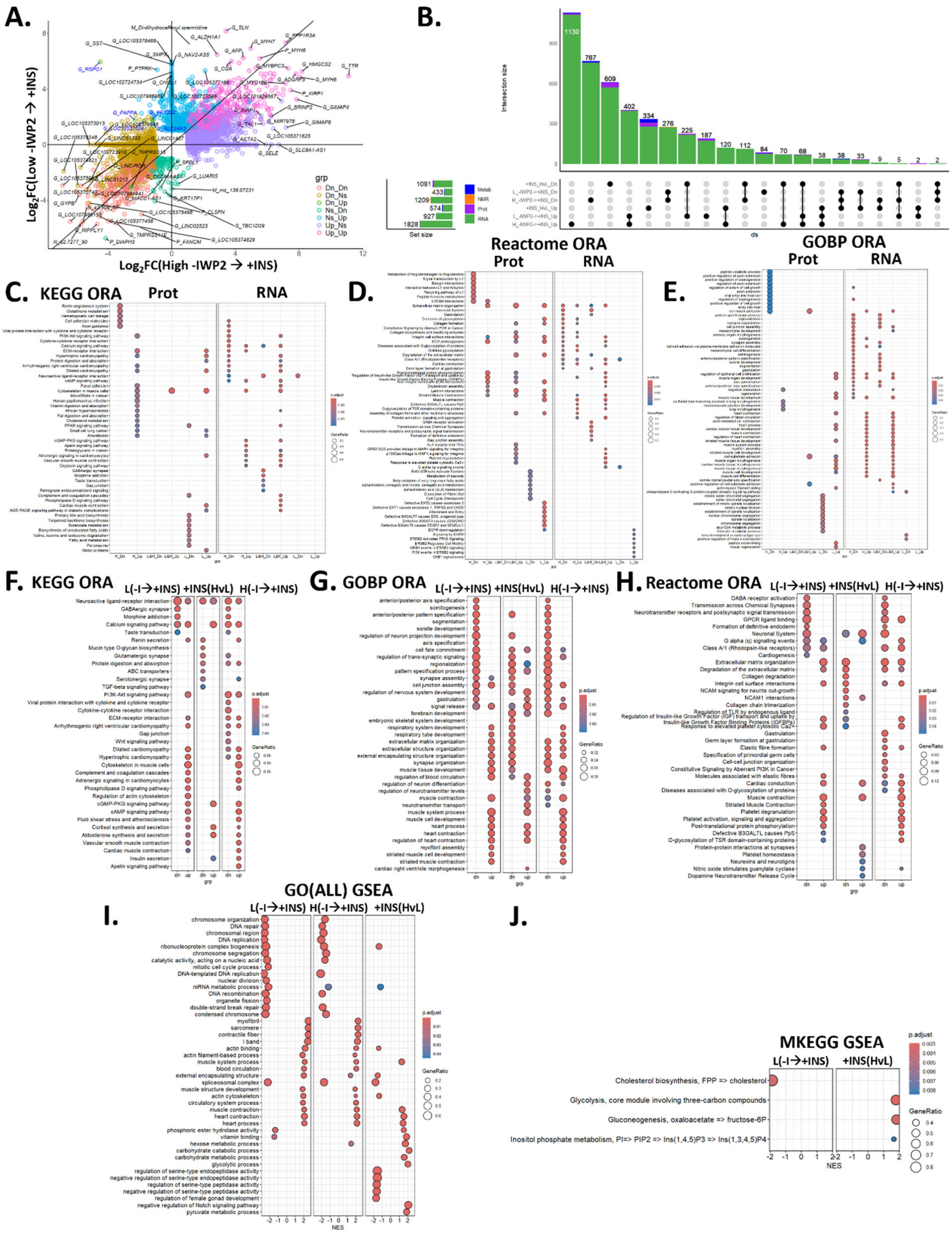


**Figure S5. Differential feature abundance, comparison of proteomic and transcriptomic pathway analysis, and additional transcriptomic pathway analysis for the -IWP2 to +INS transition. A)** Parity plot comparing the differential features changing from the -IWP2 step to the +INS step in high (x-axis) and low (y-axis) efficiency differentiations. **B)** UpSet plot of differential features for each comparison (as in Figure 4H) further separated by direction of change where Up corresponds to enriched and Dn (Down) corresponds to decreased for the temporal or high (H) versus low (L) differentiation efficiencies as indicated. **C-E)** Overrepresentation analysis for differentially expressed proteomic and transcriptomic features for the **C)** KEGG, **D)** Reactome, **E)** GO Biological Process databases. Comparisons are denoted as H (high efficiency), L (low efficiency), Up (enriched), and Dn (decreased) on the x-axis. Comparisons shown are enriched during the -IWP2 to +INS transition in either high, low, or both differentiation efficiency groups as indicated. **F-H)** Overrepresentation analysis for differentially expressed transcriptomic features for the **F)** KEGG, **G)** GO Biological Process, **H)** Reactome databases. Comparisons are shown for temporal or high (H) versus low (L) differentiation efficiencies as indicated. **I-J)** Gene set enrichment analysis for differentially expressed transcriptomic features for the **I)** GO and **J)** Metabolic KEGG databases. Comparisons are shown for temporal or high (H) versus low (L) differentiation efficiencies as indicated.

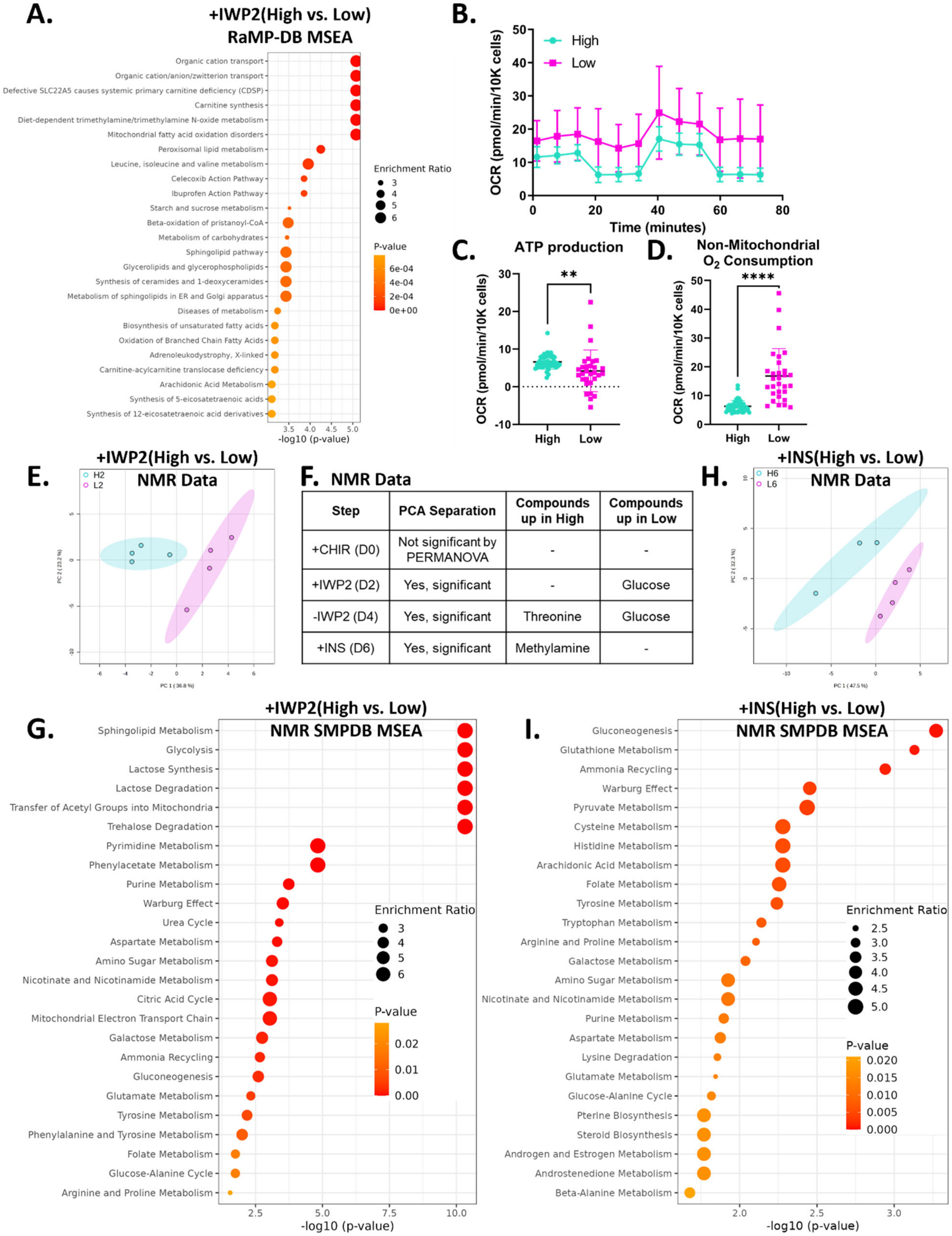


**Figure S6. Intracellular and extracellular metabolomics and oxygen consumption rate metrics distinguish high and low efficiency hPSC-CM progenitors. A)** MSEA for differentially abundant intracellular metabolites for the RaMP-DB database displaying the top 25 pathways enriched in high efficiency progenitors at the +IWP2 differentiation step (FDR adjusted p-value < 0.05, raw p-value displayed). **B-D)** MitoStress test at the +INS differentiation step (Day 6) for cryopreserved cardiac progenitor cells from high (turquoise, n = 46 well replicates) and low (magenta, n = 30 well replicates) efficiency hPSC-CM differentiations in WTC11 hiPSCs. P-values from unpaired t-tests. **B)** Representative traces of the real-time oxygen consumption rate (OCR) for high and low efficiency cardiac progenitors at the +INS differentiation step (Day 6). **C-D)** Analysis of ATP-linked respiration and **D)** non-mitochondrial oxygen consumption from Seahorse MitoStress test. **E)** PCA of the NMR extracellular metabolome at the +IWP2 step (Day 2) for high (turquoise, H2) and low (magenta, L2) differentiation efficiencies (95% CI, n = 4 well replicates per group). **F)** Table of PCA separation metrics and differentially abundant NMR extracellular metabolites enriched at each early differentiation step (Log2FC > 1, FDR adjusted p-value < 0.05). **G)** MSEA for differentially abundant NMR extracellular metabolites for the SMPDB database displaying the top 25 pathways enriched in high efficiency progenitors at the +IWP2 differentiation step (FDR adjusted p-value < 0.05, raw p-value shown). **H)** PCA of the NMR extracellular metabolome at the +INS step (Day 6) for high (turquoise, H6) and low (magenta, L6) differentiation efficiencies (95% CI, n = 3-4 well replicates per group). **I)** MSEA for differentially abundant NMR extracellular metabolites for the SMPDB database displaying the top 25 pathways enriched in high efficiency progenitors at the +INS differentiation step (FDR adjusted p-value < 0.05, raw p-value shown). All data are represented as mean ± SD.

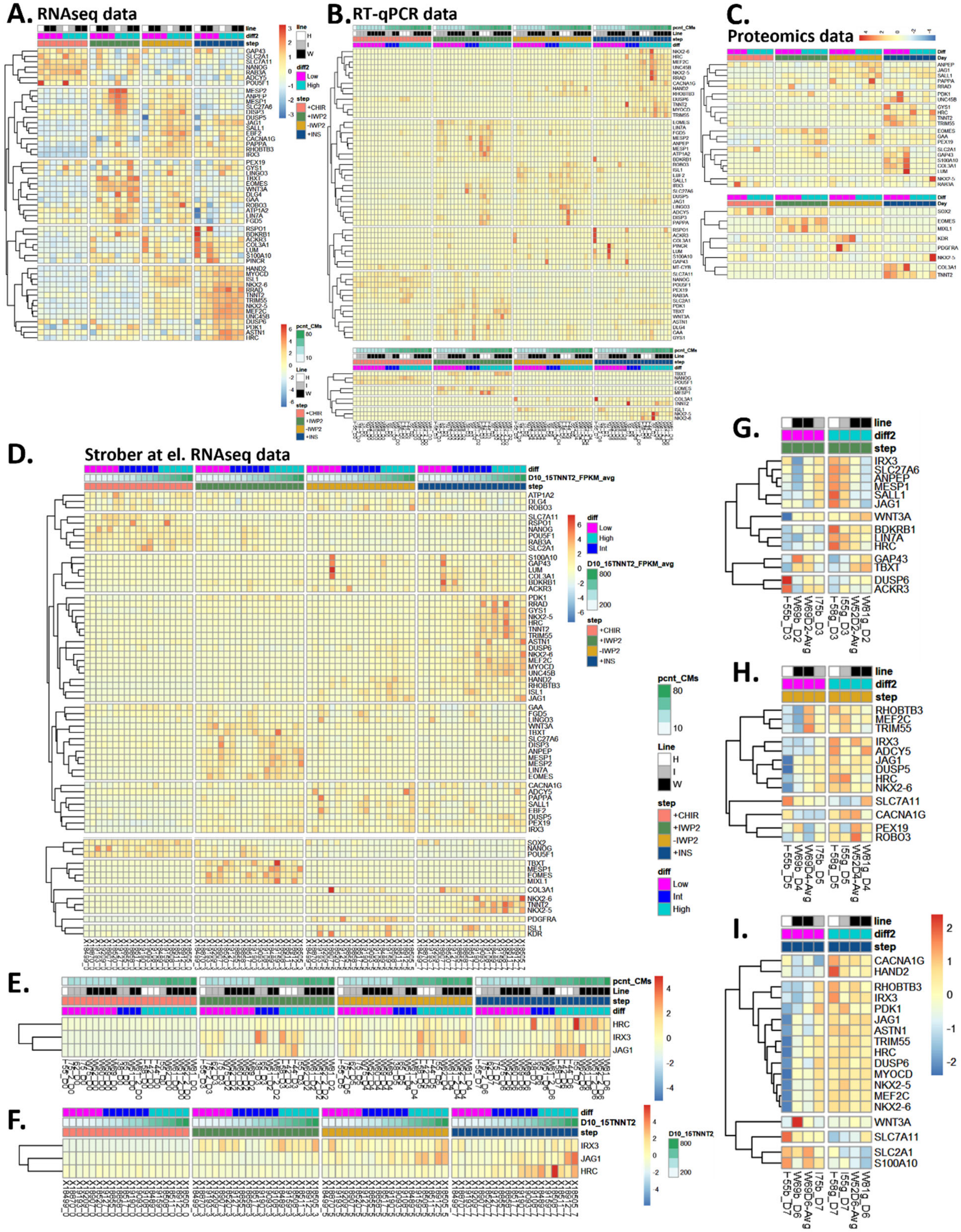


**Figure S7. Heatmaps of putative novel marker genes and canonical marker genes across transcriptomic and proteomic datasets and RT-qPCR validation. A)** Heatmap of 54 candidate marker genes differentially expressed in high versus low efficiency CM differentiations, informed from RNAseq data (TPM). **B)** Heatmap of candidate marker genes assessed in RT-qPCR samples for 22 differentiations where the top portion corresponds to the 54 candidate marker genes shown in panel A and the bottom corresponds to canonical marker genes (2^-ΔCT). **C)** Heatmap of the candidate marker genes in the proteomics data where the top portion corresponds to 11 detected markers from the top 31 and the bottom portion corresponds to the eight detected canonical markers (LFQ). **D)** Heatmap of the top marker genes identified in the Strober et al. RNAseq where the top portion corresponds to candidate marker genes and the bottom corresponds to canonical marker genes (Day 10-15 *TNNT2* FPKM average). **E-F)** Heatmaps of the expression of the three conserved predictor genes (*IRX3*, *HRC*, and *JAG1*) in the **E)** RT-qPCR data (2^-ΔCT) and **F)** the Strober et al. dataset (Day 10-15 *TNNT2* FPKM average). **G-I)** Heatmaps of the top differentiation step-specific predictor genes at the **G)** +IWP2, **H)** -IWP2, and **I)** +INS differentiation steps from RNAseq data (TPM; complementary to Figure 6E,J,N). TPM = transcripts per million. LFQ = Label free quantitation. FPKM = Fragments Per Kilobase of transcript per Million mapped reads. pcnt_CMs = percentage of CMs (flow cytometry of cTnT). ΔCT = cycle threshold of gene minus cycle threshold of the geomean of 3 reference genes (*ZNF384*, *EDF1*, *DDB1*). Line = cell line (H = H9, I = IMR90-4, W = WTC11).

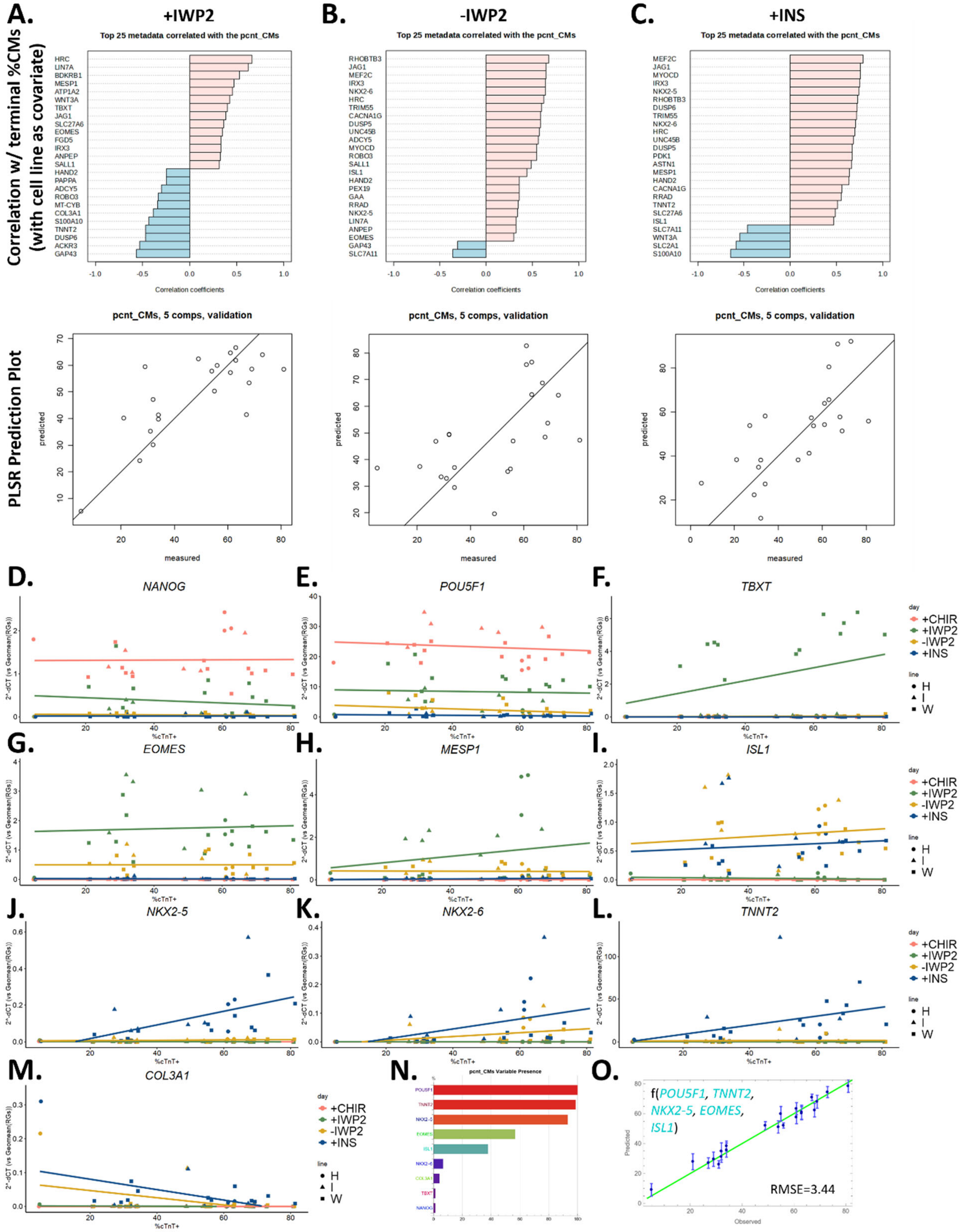

**Figure S8. Model development for prediction of terminal cardiomyocyte purity from RT-qPCR data using linear models and canonical marker genes. A-C)** Linear model development at the **A)** +IWP2, **B)** -IWP2, and **C)** +INS differentiation steps using RT-qPCR data to predict cardiomyocyte purity (2^-ΔCT). Linear regression of the top 25 predictor genes with the terminal percentage of cTnT+ cells (top row). Partial Least Squares Regression (PLSR) prediction plots based on the step-specific predictor genes presented in Figure 6E,J,N (bottom row). **D-M)** Linear regression plots of canonical marker gene expression (RT-qPCR 2^-ΔCT vs terminal %cTnT+) for each differentiation step. **N-O)** Non-linear model prediction of the percentage of cTnT+ cells from 10 canonical markers genes including *TNNT2* at the +INS differentiation step. **N)** Variable presence in evolved models after one round of predictive modeling for 10 canonical cardiomyocyte differentiation markers. **J)** Nonlinear prediction for the top 5 canonical marker genes after one round model evolution. The predictive function is listed as f(*genes*) with turquoise genes enriched in high efficiency differentiations and magenta genes enriched in low efficiency differentiations. pcnt_CM = percentage of CMs (flow cytometry of cTnT). ΔCT = cycle threshold of gene minus cycle threshold of the geometric mean of 3 reference genes (*ZNF384*, *EDF1*, *DDB1*). RMSE = root mean squared error.

**Table S1.** qPCR targets with forward and reverse primers and amplicon sizes in base pairs.

| Target | FW Primer | RV Primer | Amplicon Size |
|---|---|---|---|
| *ZNF384* | AATCTGCAGTCCCACAGACG | ACTGTGTGCGTAGACAGGTG | 116 |
| *EDF1* | CCAAGCAGGCTATCTTAGCGG | GACCTTGCTGGATCACCTTG | 183 |
| *DDB1* | TCAACGGCATGATAGGGCTG | CGCTCGGTGTGAAAGGATCT | 142 |
| *NANOG* | CGAAGAATAGCAATGGTGTGACG | TTCCAAAGCAGCCTCCAAGTC | 329 |
| *POU5F1* | CAGTGCCCGAAACCCACAC | GGAGACCCAGCAGCCTCAAA | 161 |
| *TBXT* | AAGAAGGAAATGCAGCCTCA | TACTGCAGGTGTGAGCAAGG | 101 |
| *MESP1* | CGTCCCCGCTCCTTCCG | GTCCCTTGTCACTTGGGCTC | 100 |
| *EOMES* | AGCCCTCAAAGACCCAGACT | TGAAGCGGTGTACATGGAATCAT | 152 |
| *ISL1* | CCTGCTTTTCAGCAACTGGTCA | AGGACTGGCTACCATGCTGT | 123 |
| *NKX2-6* | CCTGAGAATGGACGCAGAGC | TTCAGGCCGGGCTGTGG | 159 |
| *NKX2-5* | CTAGAGCCGAAAAGAAAGAGCTG | CGCCGCTCCAGCTCATAGA | 151 |
| *TNNT2* | TTCACCAAAGATCTGCTCCTCGCT | TTATTACTGGTGTGGAGTGGGTGTGG | 165 |
| *COL3A1* | TCAGGATCCGTTCTCTGCGA | AGGGCGAGTAGGAGCAGTTG | 126 |
| *HRC* | AACATGGGTGAGCACTGCG | TGTCTGCCAGGGCCTGATAAAG | 136 |
| *MEF2C* | TGACTGTGAGATTGCGCTGA | CTTCTTTCTCAACGTCTCCACG | 157 |
| *UNC45B* | ATCCGCACACTGGTGGGAC | GCCACTTGCGACACTGTTTG | 115 |
| *RRAD* | CGACTCAGACGAGAGCGTTT | GATCATAGGTGTGCCCTGCT | 131 |
| *CACNA1G* | TAAATCGGACTGTGCCGAGG | TTGTGGTTCATGATGGGCTGCT | 177 |
| *RHOBTB3* | CTAGCTGACCACCTTGGGTT | TGTCCAACGACTCGGACATC | 121 |
| *DUSP6* | CCTCTACTTGGGCTGTGCCA | AAGACACCACAGTTCTTGCCC | 231 |
| *MYOCD* | GGGTTGTTAGCTGCGGTCAG | CTGGACGTTTCAGTGGTGGT | 234 |
| *TRIM55* | CACACTTGGGGACAGCGAG | AATACGGGTTAGAGGCCTGGAAAA | 206 |
| *LIN7A* | TTGCAGCTAGTGAAGGCCAC | TCTCCTTCCACACTCACTCCG | 208 |
| *FGD5* | CGGGCCTGATGAGAGAGCG | TTACACCGGCTGAGATAGCCA | 199 |
| *MESP2* | CGCAGTGTACCAGGGTCTCTC | CTGGTCCTCTGAGTTGGGCA | 157 |
| *ANPEP* | TCAGGCAATGAGTGGGTCCT | AGGGACCTTATGGGCACTGG | 183 |
| *MESP1* | CGTCCCCGCTCCTTCCG | GTCCCTTGTCACTTGGGCTC | 100 |
| *ATP1A2* | GCTAAGGTCCCTCAGCCACT | GCTTGTGGTCATCCATTGCC | 160 |
| *BDKRB1* | CCGTCGGCCCAGACTGAAG | CCAGGATGATGCCATGCACAG | 135 |
| *ROBO3* | GTCCGTACACAGGATAGCAGC | CTCCAAGACACCCGGAAACC | 182 |
| *EBF2* | GCACAGTGGCATGATGGGAA | GTGTTGCGGATGTACCCTTGA | 99 |
| *SALL1* | CATCGAGTGCCCTGCAGATT | CACTTCCTGATCTTGCCGCC | 249 |
| *IRX3* | GGCGGAACAGATCGCTGTAG | AGAGCCGATAAGACCAGGGC | 119/315 |
| *SLC27A6* | GTGTTGAGTTGGGTGCCACT | CCAATTGCCAAACGCACCTT | 178 |
| *DUSP5* | CAGCCCTGCTGAATGTCTCC | CCTTTTCCCTGACACAGTCAATG | 150 |
| *JAG1* | GCCAAGTGCCAGGAAGTTTCA | CGATCCGTCCATTCAGGCAC | 115 |
| *LINGO3* | GAGGGCTGAAGTTCGTGCAA | AACCCTAACCTGGCCGAGAA | 180 |
| *ADCY5* | CTCTGAGGGCATCTGGTGGA | GAGAACATTGGAGACAAGCTGC | 178 |
| *DISP3* | TGCCCTCCTTCCAGGTGTAT | CATCACAGGCCAAGGTCGTC | 135 |
| *PAPPA* | CCAGCCGGTGCTATTTCCAT | TGGTTCGACACACCTGGGAT | 201 |
| *RSPO1* | CGACATGAACAAGTGCATCAAATG | ACGGAGACCACTCGCTCAT | 200 |

| | | | |
|---|---|---|---|
| *ACKR3* | TAAGGGAGCCAGCGCACAG | GCAGATCCATCGTTCTGAGGC | 111 |
| *PINCR* | ACTGACTGTGCAAGAACCCT | AGAGACCAGAAAGACAGCAGGT | 90 |
| *LUM* | TGGTTGAGCTGGATCTGTCCT | CGATTGCCATCCAAACGCAAA | 178 |
| *S100A10* | ATGGAACACGCCATGGAAAC | TCCACAGCCAGAGGGTCTTTT | 146 |
| *GAP43* | CCACTGATAACTCGCCGTCC | CCGGGCACTTTCCTTAGGTT | 227 |
| *MT-CYB* | CTTCCTACACATCGGGCGAG | GTGTGAGGGTGGGACTGTCT | 248 |
| *SLC7A11* | TGCATCGTCCTTTCAAGGTGC | CACCTGGGTTTCTTGTCCCA | 178 |
| *PEX19* | AAATGCCACTGACCTTCAGAACT | TCCCGATGACTCTGCAACCA | 210 |
| *RAB3A* | ATCAAGCTGCAGATCTGGGACAC | CTGATGACACCACCCGCTCA | 232 |
| *SLC2A1* | GGCTTCTCCAACTGGACCTC | CCGGAAGCGATCTCATCGAA | 176 |
| *PDK1* | ACCTCGTGTTGAGACCTCCC | GCATCTGTCCCGTAACCCTCT | 132 |
| *WNT3A* | CCGCTTCTGCAGGAACTACG | CCGACTCCCTGGTAGCTTTG | 170 |
| *ASTN1* | AGACACCTATCCTGGACGGC | GGCATTGTCACTTCCTGGTGT | 176 |
| *DLG4* | TATGAGTTGCAGGTGAACGGG | TGTCGTTGACCCTGAGGCG | 199 |
| *GAA* | GACGTCCAGTGGAACGACCT | ATACCTTCCCAATCAGCGGC | 247 |
| *GYS1* | ATTCGACCTGGAGAACGCAG | CGAAATACACCTTGCAGCCC | 248 |
| *nucDNA* | CAACTTCATCCACGTTCACC | GAAGAGCCAAGGACAGGTAC | |
| *mitoDNA* | CGAAAGGACAAGAGAAATAAGG | CTGTAAAGTTTTAAGTTTTATGCG | |

**Table S2.** RT-qPCR differentiation metadata and DataModeler predictions for cardiomyocyte purity (pcnt_CMs) for each differentiation step and for all step-specific predictor genes versus canonical genes.

| Diff Metadata | | +IWP2_AllModelGenes | | +IWP2_CanonicalModelGenes | | -IWP2_AllModelGenes | | -IWP2_CanonicalModelGenes | | +INS_AllModelGenes | | +INS_CanonicalModelGenes_NO_TNNT2 | | +INS_AllCanonicalModelGenes | |
|---|---|---|---|---|---|---|---|---|---|---|---|---|---|---|---|
| Diff# | pcnt_CMs | Prediction | Error | Prediction | Error | Prediction | Error | Prediction | Error | Prediction | Error | Prediction | Error | Prediction | Error |
| 1 | 5 | 2.951624 | -2.04838 | 8.069763 | 3.069763 | 4.449357 | -0.55064 | 4.679542 | -0.32046 | 5.076555 | 0.076555 | 5.137382 | 0.137382 | 8.99754 | 3.99754 |
| 2 | 21 | 22.816 | 1.815998 | 36.14476 | 15.14476 | 30.01271 | 9.012707 | 25.22496 | 4.224955 | 29.03852 | 8.038517 | 31.5228 | 10.5228 | 27.83812 | 6.838118 |
| 3 | 27 | 32.13422 | 5.134223 | 39.25218 | 12.25218 | 34.88732 | 7.887319 | 40.3957 | 13.3957 | 39.05588 | 12.05588 | 27.28053 | 0.280527 | 26.99997 | -3.30E-05 |
| 4 | 29 | 29.65481 | 0.654806 | 28.26537 | -0.73463 | 25.25043 | -3.74957 | 35.76567 | 6.76567 | 26.8452 | -2.1548 | 25.00142 | -3.99858 | 29.51471 | 0.514711 |
| 5 | 31 | 30.42779 | -0.57221 | 38.57479 | 7.574785 | 31.93243 | 0.932429 | 40.95373 | 9.953725 | 36.62899 | 5.628985 | 32.81621 | 1.816214 | 26.06472 | -4.93528 |
| 6 | 32 | 42.87971 | 10.87971 | 41.15477 | 9.154766 | 28.8983 | -3.1017 | 39.95049 | 7.950486 | 38.2613 | 6.261299 | 38.45576 | 6.455763 | 34.78832 | 2.788318 |
| 7 | 32 | 32.39935 | 0.399352 | 38.2758 | 6.275798 | 32.80337 | 0.803367 | 37.22753 | 5.227534 | 26.80898 | -5.19102 | 29.8436 | -2.1564 | 31.15165 | -0.84835 |
| 8 | 34 | 29.46787 | -4.53213 | 39.20199 | 5.201987 | 40.75089 | 6.750895 | 25.10025 | -8.89975 | 33.25799 | -0.74201 | 40.19172 | 6.191722 | 38.33561 | 4.335612 |
| 9 | 34 | 33.04598 | -0.95402 | 29.23102 | -4.76898 | 33.41818 | -0.58182 | 34.62488 | 0.62488 | 32.93441 | -1.06559 | 33.85653 | -0.14347 | 35.67619 | 1.676188 |
| 10 | 49 | 56.7781 | 7.778103 | 42.50924 | -6.49076 | 48.19345 | -0.80655 | 50.30078 | 1.300782 | 48.24102 | -0.75898 | 43.08543 | -5.91457 | 51.99034 | 2.990338 |
| 11 | 54 | 53.21257 | -0.78743 | 55.66275 | 1.662754 | 48.28596 | -5.71404 | 45.48254 | -8.51746 | 46.49193 | -7.50807 | 45.95819 | -8.04181 | 51.13501 | -2.86499 |
| 12 | 55 | 52.08081 | -2.91919 | 51.75871 | -3.24129 | 51.69685 | -3.30315 | 40.08194 | -14.9181 | 55.41223 | 0.412233 | 58.81072 | 3.810721 | 59.9221 | 4.922101 |
| 13 | 56 | 58.58823 | 2.588231 | 43.55612 | -12.4439 | 60.48672 | 4.486721 | 51.87739 | -4.12261 | 56.8203 | 0.8203 | 52.14612 | -3.85388 | 52.0861 | -3.9139 |
| 14 | 61 | 55.67205 | -5.32795 | 49.28044 | -11.7196 | 62.67811 | 1.67811 | 66.10493 | 5.104933 | 63.97757 | 2.977574 | 68.18765 | 7.187654 | 63.36373 | 2.363726 |
| 15 | 61 | 62.7664 | 1.766404 | 66.09289 | 5.092893 | 56.6115 | -4.3885 | 61.34511 | 0.345108 | 56.42185 | -4.57815 | 59.08005 | -1.91995 | 57.55857 | -3.44143 |
| 16 | 63 | 63.01462 | 0.014619 | 61.3086 | -1.6914 | 72.24766 | 9.247656 | 57.68408 | -5.31592 | 64.32156 | 1.321564 | 62.92322 | -0.07678 | 60.98268 | -2.01732 |
| 17 | 63 | 61.14064 | -1.85936 | 69.57672 | 6.576719 | 66.39015 | 3.390147 | 60.54539 | -2.45461 | 52.80074 | -10.1993 | 58.19684 | -4.80316 | 60.31946 | -2.68054 |
| 18 | 67 | 54.13502 | -12.865 | 52.01787 | -14.9821 | 59.8004 | -7.1996 | 62.52785 | -4.47215 | 71.16814 | 4.168137 | 68.30703 | 1.307029 | 70.91565 | 3.915654 |
| 19 | 68 | 66.00465 | -1.99535 | 65.44953 | -2.55047 | 58.5181 | -9.4819 | 63.37188 | -4.62812 | 61.09621 | -6.90379 | 60.88973 | -7.11027 | 62.35259 | -5.64741 |
| 20 | 69 | 65.70816 | -3.29184 | 55.07219 | -13.9278 | 78.29746 | 9.297457 | 61.28262 | -7.71738 | 68.40979 | -0.59021 | 65.49169 | -3.50831 | 67.94269 | -1.05731 |
| 21 | 73 | 73.55751 | 0.557506 | 73.73353 | 0.733529 | 70.716 | -2.284 | 79.54435 | 6.544352 | 76.71331 | 3.713311 | 72.03491 | -0.96509 | 74.46033 | 1.460325 |
| 22 | 81 | 84.14788 | 3.147876 | 79.82303 | -1.17697 | 66.9158 | -14.0842 | 79.68648 | -1.31352 | 77.0823 | -3.9177 | 73.16459 | -7.83541 | 78.41422 | -2.58578 |

**Table S3.** Model ensemble for all step-specific predictor genes at the +IWP2 differentiation step (+IWP2_AllModelGenes).

| | Complexity | $1-R^2$ | Vars | Function |
|---|---|---|---|---|
| **1** | 33 | 0.075 | ACKR3 LIN7A MESP1 SALL1 | $47.01-\frac{695.40}{7.84+\frac{\mathrm{LIN7A}}{\mathrm{ACKR3}}}-\frac{5.27}{\mathrm{MESP1}}+\frac{94.68}{\mathrm{SALL1}}$ |
| **2** | 39 | 0.062 | ACKR3 LIN7A MESP1 SALL1 | $67.84-228.52\sqrt{\mathrm{ACKR3}}-\frac{4.78}{\mathrm{LIN7A}}-\frac{4.80}{\mathrm{MESP1}}+\frac{91.87}{\mathrm{SALL1}}$ |
| **3** | 41 | 0.060 | ACKR3 LIN7A MESP1 SALL1 | $60.74-\frac{3715.86}{45.57+\frac{2}{\mathrm{ACKR3}}}-\frac{4.98}{\mathrm{LIN7A}}-\frac{4.78}{\mathrm{MESP1}}+\frac{92.27}{\mathrm{SALL1}}$ |
| **4** | 44 | 0.074 | ACKR3 LIN7A MESP1 SALL1 | $42.80-\frac{791.70}{9.56+\frac{\mathrm{LIN7A}}{\mathrm{ACKR3}}}-\frac{5.33}{\mathrm{MESP1}}+\frac{102.65}{\mathrm{SALL1}}+0.31\,\mathrm{MESP1}\,\mathrm{SALL1}$ |
| **5** | 45 | 0.059 | ACKR3 LIN7A MESP1 SALL1 | $72.57-228.51\sqrt{\mathrm{ACKR3}}-\frac{4.85}{\mathrm{MESP1}}-\frac{7.39}{\mathrm{LIN7A}+\frac{\mathrm{LIN7A}}{\mathrm{SALL1}}}+\frac{83.27}{\mathrm{SALL1}}$ |
| **6** | 45 | 0.072 | ACKR3 LIN7A MESP1 SALL1 | $-216.82-233.10\sqrt{\mathrm{ACKR3}}-61.65\,\mathrm{LIN7A}-\frac{4.17}{\mathrm{MESP1}}+\frac{21909.49}{58.43+\frac{\mathrm{SALL1}}{\mathrm{LIN7A}}}$ |
| **7** | 45 | 0.093 | ACKR3 LIN7A MESP1 SALL1 | $66.99-257.03\sqrt{\mathrm{ACKR3}}-\frac{4.11}{\mathrm{MESP1}}-\frac{8.27}{\mathrm{LIN7A}+\frac{\mathrm{MESP1}}{\mathrm{SALL1}}}+\frac{94.09}{\mathrm{SALL1}}$ |
| **8** | 46 | 0.097 | ACKR3 LIN7A MESP1 SALL1 | $118.84-241.82\sqrt{\mathrm{ACKR3}}+\frac{5.61}{\mathrm{LIN7A}}-\frac{4.19}{\mathrm{MESP1}}-\frac{5.64\,\mathrm{SALL1}}{\mathrm{LIN7A}}$ |
| **9** | 49 | 0.056 | ACKR3 LIN7A MESP1 SALL1 | $-783.60-230.99\sqrt{\mathrm{ACKR3}}-\frac{4.84}{\mathrm{MESP1}}+\frac{194865.59}{226.27+\frac{\sqrt{\mathrm{SALL1}}}{\mathrm{LIN7A}}}+\frac{77.05}{\mathrm{SALL1}}$ |

**Table S4.** Model ensemble for canonical genes at the +IWP2 differentiation step (+IWP2_CanonicalModelGenes).

| | Complexity | $1-R^2$ | Vars | Function |
|---|---|---|---|---|
| **1** | 42 | 0.216 | COL3A1<br>MESP1<br>NKX2–5<br>TBXT | $-1.98 + \frac{3.68\times10^{-6}}{\text{COL3A1}} + 24.78\sqrt{\text{MESP1}} + \frac{5.67\times10^{-6}}{\text{NKX2-5}} + 24.02\,\text{MESP1}\,\text{TBXT}$ |
| **2** | 42 | 0.246 | COL3A1<br>MESP1<br>NKX2–5<br>TBXT | $-0.38 + \frac{3.81\times10^{-6}}{\text{COL3A1}} + \frac{54.87}{1+\frac{1}{\text{MESP1}}} + \frac{5.71\times10^{-6}}{\text{NKX2-5}} + 23.66\,\text{MESP1}\,\text{TBXT}$ |
| **3** | 44 | 0.209 | COL3A1<br>MESP1<br>NKX2–5<br>TBXT | $4.91 + \frac{3.55\times10^{-6}}{\text{COL3A1}} + \frac{109.58}{1.00+\frac{6}{\text{MESP1}}} + \frac{5.57\times10^{-6}}{\text{NKX2-5}} + 24.95\,\text{MESP1}\,\text{TBXT}$ |
| **4** | 46 | 0.217 | COL3A1<br>MESP1<br>NKX2–5<br>TBXT | $-8.94 + \frac{3.15\times10^{-6}}{\text{COL3A1}} + 31.46\sqrt{\text{MESP1}} + \frac{5.11\times10^{-6}}{\text{NKX2-5}} + 14.75\sqrt{\text{MESP1}}\,\text{TBXT}$ |
| **5** | 51 | 0.197 | MESP1<br>NANOG<br>NKX2–5<br>TBXT | $78.66 - 64.97\sqrt{\text{NANOG}} + \frac{2.77\times10^{-6}}{\text{NKX2-5}} - \frac{24.21}{\text{MESP1}+\text{NANOG}+\text{NKX2-5}} + 11.67\,\text{TBXT}$ |
| **6** | 55 | 0.190 | MESP1<br>NANOG<br>NKX2–5<br>TBXT | $81.97 - 68.96\sqrt{\text{NANOG}} - \frac{25.25}{\text{MESP1}+\text{NANOG}} + \frac{4003.37}{\text{NKX2-5}+\frac{1}{\text{TBXT}}} - 3988.55\,\text{TBXT}$ |
| **7** | 62 | 0.205 | COL3A1<br>MESP1<br>NKX2–5<br>TBXT | $23.95 + 7.26\,\text{MESP1} + \frac{4.42\times10^{-6}}{\text{NKX2-5}} + 23.46\,\text{MESP1}\,\text{TBXT} - \frac{549624.46}{\text{MESP1}+\frac{\frac{2}{\text{TBXT}}+\text{TBXT}}{\text{COL3A1}}}$ |
| **8** | 62 | 0.221 | COL3A1<br>MESP1<br>NKX2–5<br>TBXT | $-1.46 + (7.45\times10^{-13})\left(58.77 + \frac{1}{\text{COL3A1}}\right)^2 + 25.53\sqrt{\text{MESP1}} + \frac{5.47\times10^{-6}}{\text{NKX2-5}} + 24.90\,\text{MESP1}\,\text{TBXT}$ |
| **9** | 78 | 0.205 | MESP1<br>NANOG<br>NKX2–5<br>TBXT | $81.90 - 73.00\sqrt{\text{NANOG}} - \frac{45.69}{2\,\text{MESP1}+\text{NANOG}+\text{NANOG}^2+\text{NKX2-5}} + \frac{4386.85}{\text{NKX2-5}+\frac{1}{\text{TBXT}}} - 4370.66\,\text{TBXT}$ |
| **10** | 100 | 0.170 | MESP1<br>NANOG<br>NKX2–5<br>TBXT | $69.25 - 44.48\,\text{NANOG} - \frac{63.92}{\sqrt{\text{MESP1}}+\text{MESP1}+\text{NANOG}+\text{NANOG}^2+\text{NKX2-5}} + \frac{5.15\times10^{-6}}{\text{NKX2-5}+10\,\text{NANOG}^4\,\text{NKX2-5}} + 10.96\,\text{TBXT}$ |

`**Table S5.** Model ensemble for all step-specific predictor genes at the -IWP2 differentiation step (-IWP2_AllModelGenes).

| | Complexity | $1-R^2$ | Vars | Function |
|---|---|---|---|---|
| 1 | 49 | 0.136 | DUSP5<br>JAG1<br>RHOBTB3<br>ROBO3 | $131.74-\frac{5.05}{\text{JAG1}}-2.01\,\text{RHOBTB3}^2-\frac{442.44}{\frac{0.86}{\text{DUSP5}}+\text{RHOBTB3}^2}-\frac{1.09\times10^{-5}}{\text{ROBO3}}$ |
| 2 | 58 | 0.121 | DUSP5<br>JAG1<br>RHOBTB3<br>ROBO3 | $107.20-\frac{5.83}{\text{JAG1}}-\frac{328.27}{\text{RHOBTB3}+\frac{\text{RHOBTB3}}{\text{DUSP5}}}-\frac{47740.54}{\text{RHOBTB3}+\frac{\text{RHOBTB3}^2}{\text{ROBO3}}}-\frac{(7.26\times10^{-5})\text{DUSP5}}{\text{ROBO3}}$ |
| 3 | 59 | 0.117 | DUSP5<br>JAG1<br>RHOBTB3<br>ROBO3 | $113.58-\frac{6.32}{\text{JAG1}}-\frac{279.24\,\text{DUSP5}}{\text{RHOBTB3}}-\frac{49023.95}{-4.41+\frac{\text{RHOBTB3}^2}{\text{ROBO3}}}-\frac{(8.50\times10^{-5})\text{DUSP5}}{\text{ROBO3}}$ |
| 4 | 62 | 0.110 | DUSP5<br>JAG1<br>RHOBTB3<br>ROBO3 | $111.97-\frac{6.14}{\text{JAG1}}-\frac{283.41}{\text{JAG1}^2+\frac{\text{RHOBTB3}}{\text{DUSP5}}}-\frac{58494.04}{\text{RHOBTB3}+\frac{\text{RHOBTB3}^2}{\text{ROBO3}}}-\frac{(8.19\times10^{-5})\text{DUSP5}}{\text{ROBO3}}$ |
| 5 | 65 | 0.116 | DUSP5<br>JAG1<br>RHOBTB3<br>ROBO3 | $114.12-\frac{6.47}{\text{JAG1}}-\frac{283.40\,\text{DUSP5}}{\text{RHOBTB3}}-\frac{(4.11\times10^{-4})\text{DUSP5}^2}{\text{ROBO3}}-\frac{47550.34\,\text{ROBO3}}{\text{RHOBTB3}^2}$ |
| 6 | 66 | 0.096 | DUSP5<br>JAG1<br>RHOBTB3<br>ROBO3 | $110.28-\frac{6.10}{\text{JAG1}}-\frac{86394.19}{\text{RHOBTB3}\left(\text{RHOBTB3}+\frac{\text{RHOBTB3}^2}{\text{ROBO3}}\right)}-\frac{(8.15\times10^{-5})\text{DUSP5}}{\text{ROBO3}}-\frac{263.61}{\frac{\text{RHOBTB3}}{\text{DUSP5}}+\text{ROBO3}}$ |
| 7 | 67 | 0.083 | DUSP5<br>JAG1<br>RHOBTB3<br>ROBO3 | $482.70-\frac{6.04}{\text{JAG1}}+\frac{629.30}{-1.69+\frac{\text{DUSP5}}{\text{RHOBTB3}}}-\frac{(8.59\times10^{-5})\text{DUSP5}}{\text{ROBO3}}-\frac{40688.17\,\text{ROBO3}}{\text{JAG1}\,\text{RHOBTB3}^2}$ |
| 8 | 70 | 0.122 | DUSP5<br>JAG1<br>RHOBTB3<br>ROBO3 | $100.05-\frac{3.11}{\text{JAG1}}-\frac{305.19}{\text{RHOBTB3}+\frac{\text{RHOBTB3}}{\text{DUSP5}}}-\frac{21597.32}{\text{JAG1}^2\left(\text{DUSP5}+\frac{\text{RHOBTB3}^2}{\text{ROBO3}}\right)}-\frac{1.46\times10^{-5}}{\text{ROBO3}}$ |
| 9 | 71 | 0.148 | DUSP5<br>JAG1<br>RHOBTB3<br>ROBO3 | $115.13-\frac{5.13}{\text{JAG1}}-\frac{326.88\,\text{DUSP5}}{\text{RHOBTB3}}-\frac{(5.89\times10^{-6})\text{RHOBTB3}}{\text{ROBO3}}-\frac{12320.32\,\text{JAG1}\,\text{ROBO3}}{\text{DUSP5}\,\text{RHOBTB3}^2}$ |
| 10 | 72 | 0.110 | DUSP5<br>JAG1<br>RHOBTB3<br>ROBO3 | $117.74-\frac{6.52}{\text{JAG1}}-\frac{303.06\,\text{DUSP5}}{\text{RHOBTB3}}-\frac{(8.89\times10^{-5})\text{DUSP5}}{\text{ROBO3}}-\frac{8980.81\left(\frac{1}{\text{DUSP5}}-\text{DUSP5}\right)\text{ROBO3}}{\text{RHOBTB3}}$ |
| 11 | 75 | 0.108 | DUSP5<br>JAG1<br>RHOBTB3<br>ROBO3 | $114.08-\frac{6.33}{\text{JAG1}}-\frac{251.89\,\text{DUSP5}}{\text{RHOBTB3}}-\frac{(9.11\times10^{-5})\text{DUSP5}}{\text{ROBO3}}-\frac{1574.06\,\sqrt{\text{RHOBTB3}\,\text{ROBO3}}}{\text{RHOBTB3}^2}$ |
| 12 | 85 | 0.127 | DUSP5<br>JAG1<br>RHOBTB3<br>ROBO3 | $100.63-56.85\,\text{DUSP5}-\frac{5.39}{\text{JAG1}}-\frac{184.60}{2+\text{RHOBTB3}+\left(\frac{1}{\text{JAG1}}-\text{RHOBTB3}^2\right)^2}-\frac{(6.35\times10^{-5})\text{DUSP5}}{\text{ROBO3}}$ |

**Table S6.** Model ensemble for canonical genes at the -IWP2 differentiation step (-IWP2_CanonicalModelGenes).

| | Complexity | $1-R^2$ | Vars | Function |
|---|---|---|---|---|
| 1 | 41 | 0.181 | ISL1<br>NKX2−6<br>POU5F1<br>TBXT | $110.49-\frac{0.72}{\text{ISL1}}-31.30\,\text{ISL1}-7.53\,\text{POU5F1}-\frac{0.55}{\frac{\text{NKX2-6}}{\text{ISL1}}+\text{TBXT}}$ |
| 2 | 45 | 0.163 | ISL1<br>NKX2−6<br>POU5F1<br>TBXT | $-0.35+\frac{35.79}{\frac{7.88\times10^{-2}}{\text{ISL1}}+\text{ISL1}}+324.63\,\text{NKX2-6}-7.04\,\text{POU5F1}+108.34\sqrt{\text{TBXT}}$ |
| 3 | 52 | 0.180 | ISL1<br>NKX2−6<br>POU5F1<br>TBXT | $151.38-\frac{132.23}{1+\frac{1}{\text{ISL1}}}+\frac{1.07\times10^{-2}}{\text{NKX2-6}}-7.19\,\text{POU5F1}-\frac{3.75}{\sqrt{\text{NKX2-6}}+\text{TBXT}}$ |
| 4 | 53 | 0.139 | ISL1<br>NKX2−6<br>POU5F1<br>TBXT | $39.17+\frac{34.67}{\frac{7.88\times10^{-2}}{\text{ISL1}}+\text{ISL1}}+351.84\,\text{NKX2-6}-0.90\,\text{POU5F1}^2-\frac{2.01}{4.76\times10^{-2}+\text{TBXT}}$ |
| 5 | 58 | 0.158 | ISL1<br>NKX2−6<br>POU5F1<br>TBXT | $22.53-\frac{0.20}{\text{ISL1}}-7.31\,\text{POU5F1}+\frac{1440.43}{17+\frac{1}{\text{ISL1}}+\frac{\text{ISL1}^2}{\text{NKX2-6}+\text{TBXT}}}$ |
| 6 | 63 | 0.135 | ISL1<br>NKX2−6<br>POU5F1<br>TBXT | $37.20+\frac{34.33}{\frac{7.88\times10^{-2}}{\text{ISL1}}+\text{ISL1}}+366.53\,\text{NKX2-6}-0.90\,\text{POU5F1}^2-\frac{1.61}{\frac{1}{25+\text{ISL1}}+\text{TBXT}}$ |
| 7 | 65 | 0.131 | ISL1<br>NKX2−6<br>POU5F1<br>TBXT | $-3.31+\frac{36.59}{\frac{7.88\times10^{-2}}{\text{ISL1}}+\text{ISL1}}-0.72\,\text{POU5F1}^2+\frac{476.55}{4\,\text{POU5F1}+\frac{1}{\text{NKX2-6}+\text{TBXT}}}$ |
| 8 | 66 | 0.136 | ISL1<br>NKX2−6<br>POU5F1<br>TBXT | $26.51-\frac{1.48}{\text{POU5F1}}-7.94\,\text{POU5F1}+12.91\,\text{TBXT}+\frac{1398.79}{17+\frac{1}{\text{ISL1}}+\frac{\text{ISL1}^2}{\text{NKX2-6}+\text{TBXT}}}$ |
| 9 | 74 | 0.158 | ISL1<br>NKX2−6<br>POU5F1<br>TBXT | $94.95-57.21\,\text{ISL1}-\frac{4.13}{\text{POU5F1}}-6.79\,\text{POU5F1}+\frac{709.59}{\frac{5}{\text{ISL1}}+\frac{1}{\text{NKX2-6}+\text{ISL1}^2\,\text{TBXT}}}$ |
| 10 | 89 | 0.129 | ISL1<br>NKX2−6<br>POU5F1<br>TBXT | $36.03-2.85\,\text{ISL1}-7.61\,\text{POU5F1}-\frac{1.59}{\text{NKX2-6}+\text{POU5F1}}+\frac{919.62}{9+\frac{2}{\text{ISL1}}+\text{TBXT}+\frac{\text{ISL1}^2}{\text{NKX2-6}+\text{TBXT}}}$ |

**Table S7.** Model ensemble for all step-specific predictor genes at the +INS differentiation step (+INS_AllModelGenes).

| | Complexity | $1-R^2$ | Vars | Function |
|---|---|---|---|---|
| 1 | 54 | 0.093 | HRC<br>RHOBTB3<br>S100A10 | $404.23-\frac{0.16}{\text{HRC}}+\frac{61.17}{\text{RHOBTB3}}-49.27\,\text{RHOBTB3}-\frac{756.54}{\frac{1}{\text{RHOBTB3}}+\text{RHOBTB3}+\frac{\text{HRC}}{\text{S100A10}}}$ |
| 2 | 54 | 0.114 | HRC<br>RHOBTB3<br>S100A10 | $458.94-\frac{0.12}{\text{HRC}}-62.87\,\text{RHOBTB3}-\frac{746.00}{\frac{1}{\text{RHOBTB3}}+\text{RHOBTB3}+\frac{\text{HRC}}{\text{S100A10}}}+0.43\,\text{S100A10}$ |
| 3 | 58 | 0.083 | HRC<br>RHOBTB3<br>S100A10 | $745.25-\frac{0.17}{\text{HRC}}-8.17\,\text{HRC}-239.27\sqrt{\text{RHOBTB3}}-\frac{930.52}{\frac{1}{\text{RHOBTB3}}+\text{RHOBTB3}+\frac{\text{HRC}}{\text{S100A10}}}$ |
| 4 | 59 | 0.105 | HRC<br>RHOBTB3<br>S100A10<br>SLC2A1 | $-548.17-\frac{0.19}{\text{HRC}}+\frac{4306.94}{6-\frac{2}{\text{RHOBTB3}}}-\frac{406.43}{\text{RHOBTB3}}-\frac{77.65\,\text{S100A10}}{\text{S100A10}+\frac{\sqrt{\text{HRC}}}{\text{SLC2A1}}}$ |
| 5 | 66 | 0.107 | HRC<br>RHOBTB3<br>S100A10 | $466.97-64.19\,\text{RHOBTB3}-\frac{761.88}{\frac{1}{\text{RHOBTB3}}+\text{RHOBTB3}+\frac{\text{HRC}}{\text{S100A10}}}+0.48\,\text{S100A10}-\frac{(4.07\times10^{-5})\,\text{S100A10}}{\text{HRC}^2}$ |
| 6 | 68 | 0.075 | HRC<br>RHOBTB3<br>S100A10<br>SLC2A1 | $271.27-\frac{0.21}{\text{HRC}}-32.72\,\text{RHOBTB3}-\frac{381.50}{\frac{1}{\text{RHOBTB3}^2}+\text{RHOBTB3}}+\frac{66.29}{\text{HRC}+\frac{\text{S100A10}\,\text{SLC2A1}}{\text{HRC}}}$ |
| 7 | 73 | 0.107 | HRC<br>RHOBTB3<br>S100A10 | $296.47-\frac{0.23}{\text{HRC}}-34.72\,\text{RHOBTB3}-\frac{422.79}{\frac{1}{\text{RHOBTB3}^2}+\text{RHOBTB3}}+\frac{309.51\,\text{HRC}}{\text{RHOBTB3}\,\text{S100A10}^2}$ |
| 8 | 74 | 0.098 | HRC<br>RHOBTB3<br>S100A10<br>SLC2A1 | $382.91-\frac{0.19}{\text{HRC}}+\frac{301.89}{\text{RHOBTB3}}-\frac{854.20}{2+\text{RHOBTB3}^2}-\frac{357.33\,\text{S100A10}}{\text{S100A10}+\frac{\text{HRC}}{\text{HRC}+\text{RHOBTB3}+\text{SLC2A1}}}$ |
| 9 | 78 | 0.072 | HRC<br>RHOBTB3<br>S100A10 | $934.52-\frac{0.23}{\text{HRC}}-299.53\sqrt{\text{RHOBTB3}}-28.39\sqrt{\text{HRC}+\frac{1}{\text{S100A10}}}-\frac{1151.54}{\frac{1}{\text{RHOBTB3}}+\text{RHOBTB3}+\frac{\text{HRC}}{\text{S100A10}}}$ |
| 10 | 86 | 0.122 | HRC<br>RHOBTB3<br>S100A10<br>SLC2A1 | $111.00-\frac{0.21}{\text{HRC}}+\frac{40.63}{\text{RHOBTB3}}-\frac{211.46}{\frac{1}{\text{RHOBTB3}^2}+\text{RHOBTB3}}+\frac{18.40\left(\sqrt{\text{HRC}}+\frac{1}{\text{S100A10}}\right)}{\text{S100A10}\,\text{SLC2A1}}$ |

**Table S8.** Model ensemble for canonical genes excluding *TNNT2* at the +INS differentiation step (+INS_CanonicalModelGenes_NO_TNNT2).

| | Complexity | $1-R^2$ | Vars | Function |
|---|---|---|---|---|
| **1** | 43 | 0.141 | ISL1<br>NANOG<br>NKX2–5 | $60.04 + 460.47\,\text{NANOG} + \frac{0.65}{\frac{1.33\times10^{-2}}{\text{ISL1}}+\text{NANOG}} - \frac{760.14}{12.45+\frac{1}{\text{NANOG}\left(1+\frac{1}{\text{NKX2–5}}\right)}}$ |
| **2** | 47 | 0.114 | ISL1<br>NANOG<br>NKX2–5<br>POU5F1 | $85.21 + 215.33\,\text{NANOG} - \frac{496.05}{8+\frac{\text{NKX2–5}}{\text{NANOG}}} - \frac{3.86}{\text{ISL1}+\frac{\text{NANOG}}{\text{POU5F1}}} - \frac{1.50\times10^{-5}}{\text{POU5F1}}$ |
| **3** | 47 | 0.114 | ISL1<br>NANOG<br>NKX2–5<br>POU5F1 | $57.29 + 441.02\,\text{NANOG} + \frac{0.64}{\frac{1.33\times10^{-2}}{\text{ISL1}}+\text{NANOG}} - \frac{699.26}{12.45+\frac{\text{NKX2–5}}{\text{NANOG}}} - \frac{9.58\times10^{-6}}{\text{POU5F1}}$ |
| **4** | 48 | 0.133 | ISL1<br>MESP1<br>NANOG<br>NKX2–5 | $-7.33 + \frac{102.29}{1+\frac{1}{\text{ISL1}}} + 625.74\,\text{NANOG} - 711.93\,\text{ISL1}\,\text{NANOG} + \frac{2.98}{\text{MESP1}+\frac{\text{NANOG}}{\text{NKX2–5}}}$ |
| **5** | 49 | 0.076 | ISL1<br>NANOG<br>NKX2–5<br>POU5F1 | $23.01 + \frac{3.84}{\frac{\text{NANOG}}{\text{NKX2–5}}+\text{NKX2–5}} - \frac{3.70}{\text{ISL1}+\frac{\text{NANOG}}{\text{POU5F1}}} + \frac{227.48}{\frac{1}{\text{NKX2–5}}+\frac{\text{NKX2–5}}{\text{POU5F1}}}$ |
| **6** | 49 | 0.122 | ISL1<br>NANOG<br>NKX2–5<br>POU5F1 | $57.81 + \frac{53.09}{\frac{1}{\text{ISL1}}+\text{ISL1}} + 267.67\,\text{NANOG} - \frac{628.02}{10.59+\frac{\text{NKX2–5}}{\text{NANOG}}} - \frac{1.32\times10^{-5}}{\text{POU5F1}}$ |
| **7** | 52 | 0.105 | ISL1<br>NANOG<br>NKX2–5<br>POU5F1 | $32.77 - 12.03\,\text{ISL1} - \frac{9.26\times10^{-2}}{\text{NANOG}} + \frac{0.29\,\text{ISL1}}{\text{NANOG}} + \frac{240.05}{\frac{1}{\text{NKX2–5}}+\frac{\text{ISL1}}{\text{POU5F1}}}$ |
| **8** | 55 | 0.098 | ISL1<br>NANOG<br>NKX2–5<br>POU5F1 | $13.01 - 20.02\,\text{ISL1} + \frac{5.97}{\text{NKX2–5}+\frac{1}{\text{ISL1}\left(6+\frac{\text{NKX2–5}}{\text{NANOG}}\right)}} + \frac{285.72}{\frac{1}{\text{NKX2–5}}+\frac{\text{NKX2–5}}{\text{POU5F1}}}$ |
| **9** | 57 | 0.060 | ISL1<br>NANOG<br>NKX2–5<br>POU5F1 | $17.84 + \frac{5.28\times10^{-2}}{\text{NKX2–5}} + \frac{4.54}{\frac{\text{NANOG}}{\text{NKX2–5}}+\text{NKX2–5}} - \frac{4.57}{\text{ISL1}+\frac{\text{NANOG}}{\text{POU5F1}}} + \frac{254.43}{\frac{1}{\text{NKX2–5}}+\frac{\text{NKX2–5}}{\text{POU5F1}}}$ |

**Table S9.** Model ensemble for all canonical genes at the +INS differentiation step (+INS_AllCanonicalModelGenes).

| | Complexity | $1-R^2$ | Vars | Function |
|---|---|---|---|---|
| **1** | 36 | 0.096 | EOMES<br>NKX2–5<br>POU5F1<br>TNNT2 | $4.62 + \frac{408.32}{8.90+\frac{0.29}{\text{NKX2–5}}} - \frac{3.91\times10^{-5}}{\text{POU5F1}} + 67.56\,\text{EOMES}\,\text{TNNT2}$ |
| **2** | 44 | 0.066 | EOMES<br>NKX2–5<br>POU5F1<br>TNNT2 | $6.44 - \frac{1.62\times10^{-2}}{\text{EOMES}} - \frac{2.09\times10^{-5}}{\text{POU5F1}} + \frac{407.53}{7+\frac{\text{POU5F1}}{\text{NKX2–5}}} + 1.62\,\text{POU5F1}\,\text{TNNT2}$ |
| **3** | 49 | 0.094 | ISL1<br>NKX2–5<br>POU5F1<br>TNNT2 | $-54.04 + 43.30\,\text{ISL1} - \frac{2.68}{\sqrt{\text{NKX2–5}}+\frac{\text{NKX2–5}}{\text{POU5F1}}} - \frac{2.44\times10^{-5}}{\text{POU5F1}} + \frac{100.62}{1+\frac{1}{\text{TNNT2}}}$ |
| **4** | 50 | 0.071 | EOMES<br>NKX2–5<br>POU5F1<br>TNNT2 | $-20.48 + \frac{207.39}{5+\frac{2}{\text{POU5F1}}} - \frac{3.51\times10^{-5}}{\text{POU5F1}} + \frac{507.01}{7.93+\frac{\text{POU5F1}}{\text{NKX2–5}}} + 58.33\,\text{EOMES}\,\text{TNNT2}$ |
| **5** | 53 | 0.066 | EOMES<br>NKX2–5<br>POU5F1<br>TNNT2 | $6.47 - \frac{1.62\times10^{-2}}{\text{EOMES}} - \frac{2.79\times10^{-2}}{\sqrt{\text{POU5F1}}} + \frac{408.00}{7+\frac{\text{POU5F1}}{\text{NKX2–5}}} + 1.62\,\text{POU5F1}\,\text{TNNT2}$ |
| **6** | 56 | 0.058 | ISL1<br>NKX2–5<br>POU5F1<br>TNNT2 | $-26.11 + \frac{860.62}{14.47+\frac{\text{POU5F1}}{\text{NKX2–5}}} + 33.96\sqrt{\text{ISL1}\sqrt{\text{POU5F1}}\sqrt{\text{TNNT2}}}$ |
| **7** | 64 | 0.051 | ISL1<br>NKX2–5<br>POU5F1<br>TNNT2 | $-28.22 + \frac{916.06}{14.47+\frac{\text{POU5F1}}{\text{NKX2–5}}} + 34.98\sqrt{\text{ISL1}\sqrt{\text{POU5F1}}\sqrt{\text{TNNT2}}} - \left(6.84\times10^{-2}\right)\text{TNNT2}$ |
| **8** | 67 | 0.040 | ISL1<br>NKX2–5<br>POU5F1<br>TNNT2 | $-33.02 + \frac{988.34}{14.33+\frac{\text{POU5F1}}{\text{NKX2–5}}} + 38.40\sqrt{\text{ISL1}\sqrt{\text{POU5F1}}\sqrt{\text{TNNT2}}} - 0.38\,\text{ISL1}\,\text{TNNT2}$ |
| **9** | 69 | 0.077 | EOMES<br>NKX2–5<br>POU5F1<br>TNNT2 | $56.44 - \frac{4.66}{3.64\times10^{-2}+\sqrt{\text{NKX2–5}}} - \frac{4.17\times10^{-5}}{\text{POU5F1}} - \frac{19.32}{\text{NKX2–5}+\frac{1}{\text{POU5F1}}+\text{POU5F1}} + 69.74\,\text{EOMES}\,\text{TNNT2}$ |
